# Evaluation of *Crenobia alpina* and *Crenobia montenigrina* from Parâng Mountains (Carpathians, Romania) according to the morphological and biological concepts of species

**DOI:** 10.1101/2023.02.08.527608

**Authors:** Babalean Anda Felicia

**Affiliations:** University of Craiova, Faculty of Horticulture, Department of Biology and Environmental Engineering, 13 A.I. Cuza Street

## Abstract

This paper assesses the taxonomic status - species or subspecies of the monopharyngeal and polypharyngeal *Crenobia* in Parâng Mt. Romania. From the morphological point of view, the two forms belong to *Crenobia alpina*, respectively to *Crenobia montenigrina*. The analysis of the copulatory apparatus indicates a seasonal reproductive isolation (restriction from reproduction) of the two forms: the monopharyngeal *alpina* become able to mate during the warm season while the polypharyngeal *montenigrina* become able to mate during the cold season. Data on the reproductive biology and peculiar morphological data are also given. The study concludes that the monopharyngeal and the polypharyngeal Crenobia from Parâng Mt. are good species according to both morphological and biological concepts.

## 1. Introduction

*Crenobia alpina* (Dana, 1766) is a freshwater flatworm with a limited dispersal ability which lives almost exclusively in cold springs and headwaters of mountainous areas, distributed in Europe and Turkey (Brändle et al. 2007, Brändle et al. 2017, Sluys 2022).

Taxonomically, *Crenobia alpina* is a complex of variants (*Crenobia alpina* sensu lato) with different morphology, reproductive biology, karyology, ecological requirements, habitat, geographic distribution (Sluys, 2022). The various forms of this complex were regarded either as good species within the genus Crenobia or as subspecies of *Crenobia alpina* – see the synthetic review of Sluys (2022).

Regardless of whether they are species or subspecies, the genus Crenobia comprises two distinct morphological types in terms of the number of pharynges:

- the monopharyngeal type includes the variants *Crenobia alpina septentrionalis, Crenobia alpina meridionalis* = *Crenobia alpina alpina, Crenobia alpina alba, Crenobia alpina bathycola, Crenobia alpina corsica*
- the polypharyngeal type includes the variants *Crenobia alpina montenigrina* with 10-15 pharynxes, *Crenobia alpina anophthalma* with 3 pharynxes, *Crenobia alpina teratophila* (very similar with *montenigrina*) and a form with 24 – 35 pharynxes from Bulgaria, described by Chichkoff as *Phagocata cornuta* (Codreanu, 1956).

The forms present in the Romanian fauna are the monopharyngeal *Crenobia alpina* and the polypharyngeal *Crenobia montenigrina* (Codreanu 1956, Babalean 2020, Sluys 2022). In Parâng Mt. the two forms are sympatric and realize mixed populations (Babalean 2020).

Several species concepts have evolved during the last century – the biological, ecological, evolutionary, phylogenetic, phenetic, genotypic concepts of species (De Quieroz 2007). De Quieroz (2007) also points out the confusion between species concepts and species delimitation, of major importance in biodiversity studies, in establishing the number of species.

Bănărescu (1973) opines citing and embracing Mayr’s school that the only objective taxon is the species reproductive isolated. The species reproductive isolated is the only objective reality, being the form and way species exist and are naturally separated from other species. However, reproductive isolation of species is not the same with restriction from reproduction (for instance, populations restricted from reproduction by physical barriers – vast spaces – which in fact, are able to give viable and fertile offspring). The biological concept of species has nothing subjective. Here I embrace the biological concept of species.

Codreanu (1956) indicates a possible reproductive isolation of *C. alpina* and *C. montenigrina* in his studied area.

The aim of this paper is the evaluation of the taxonomic status (species or subspecies) of the two forms from Parâng Mt. (the monopharyngeal *alpina* and the polypharyngeal *montenigrina*) according to the morphological and biological concepts of species.

## 2. Material and method

The study is based on more than 300 specimens sampled from Parâng Mt. (Rânca Resort) between 1.07.2016 and 7.11.2022, from populations occupying different habitats (Fig.1): reocrene springs, springs with sandy reservoirs, springs with catching reservoirs, swampy area, spring collecting streams at various altitudes. The sampling was done during summer (June, July) and during the cold season (November).

**Fig. 1.**
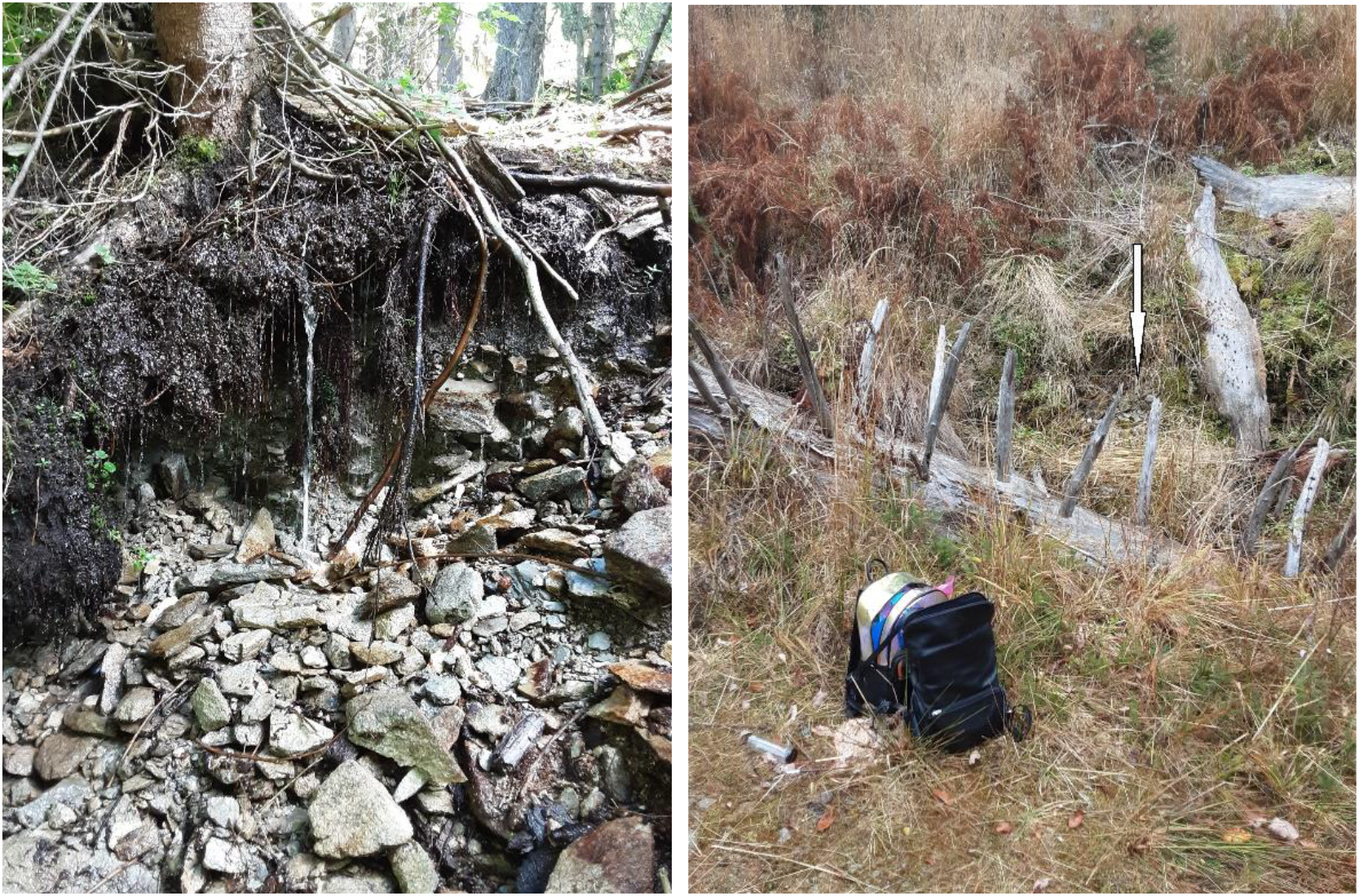
Habitates: spring no. 3 (left) and no. 2 (right)

### Sampling remark

in very few locations all the visible specimens of the population were collected, namely, only where 3 - 5 specimens were spotted (only one location, at the teleski area). For the rest of the populations, a 1/4th to a maximum a half of the visible specimens were collected. Part of the worms were fixed in Beauchamp solution for 24 hours, thereafter, removed and stored in ethanol 75º; another part of the worms was fixed and stored in 96º ethanol. The maturity of the copulatory area was assessed on histological sections and by visual inspection of fixed, uncut worms. In this latter case the specimens with large copulatory area were considered capable to realize the sperm transfer during mating. Some specimens were processed by usual techniques for Haematoxilin-Eosin staining, for histological frontal and sagittal serial sections at 5 μm:

- CR1 – monopharyngeal, 1.07.2016, Rânca (Cab. Bot. – Romanul), sagittal sections on 53 slides,
- CR2 – polypharyngeal 1.07.2016, Rânca (Cab. Bot. – Romanul), sagittal sections on 43 slides,
- CR3 – polypharyngeal, 28.06.2017, Rânca – swampy area at the triple fountain spring, frontal sections on 10 slides,
- CR4 – monopharyngeal, 1.07.2016, Rânca (Cab. Bot. – Romanul), frontal sections on 12 slides,
- CR5 – polipharyngeal, 2.07.2016, spring Paltin Mt., frontal sections requested; worm wasted in a doubtful and improbable manner,
- CR6 – monopharyngeal, 28.06.2017, Rânca, teleski area, sagittal sections requested; worm wasted in a doubtful and improbable manner,
- CR7 – monopharyngeal, 2.07. 2016, Rânca – collecting stream at Paltin Mt., frontal sections requested; worm wasted in a doubtful and improbable manner,
- CR8 – polypharyngeal, 5.11.2020, Rânca – spring with pipe, reservoir and cross at Păpuşa Mt., sagittal sections on 71 slides,
- Cm1 – monopharyngeal, 2.07.2016, Rânca - collecting stream at Paltin Mt., sagittal sections on 63 slides,
- Cm2 – monopharyngeal, 5.07.2021, Rânca (Cab. Bot. – Romanul, reocrene spring no. 1), sagittal sections requested; transversal sections on 25 slides, the worm was not entirely cut,
- Cm3 – monopharyngeal, 5.07.2021, Rânca (Cab. Bot. – Romanul, collecting stream below the pipe), sagittal sections on 28 slides (compromised, the copulatory area removed from all slides),
- Cp1 – polypharyngeal, 28.07.2017, Rânca, swampy area at the triple fountain, sagittal sections on 40 slides,
- Cp2 – polypharyngeal, 10.07.2016, Rânca – Tidvele, spring at the highest altitude (1841m), frontal sections on 9 slides.

The reconstruction of the copulatory apparatus was done by drawings on transparent tracing paper. The serial histological sections were photographed with a Levenhuk camera, the microphotographs were printed on A3 format sheets and serially drawn on transparent tracing paper.

Abbreviations used in the figures: af – atrial fold, bc – bursal canal, cb – copulatory bursa, cov – common oviduct, fa – fibrous atrium (fibrous part of the atrium), fdo – fine ducts orifices, ga – genital atrium (the distal part), go – gonopore, ov – oviduct, ph – pharynx, pp – penis papilla, vd – vas deferens.

## 3. Results

### 3.1 Types of populations

In the studied area, *Crenobia alpina* and *Crenobia montenigrina* realized the following types of populations with respect to pharyngy state (mono- / polypharyngeal) and the number of collected specimens:

#### A) Pure populations

a. exclusively monopharyngeals, with a small number of specimens – only one monopharyngeal population (teleski area)
b. large populations, exclusively polypharyngeals

#### B) mixed populations

a. predominant monopharyngeals
b. predominant polypharyngeals
c. nearly equal proportions of mono- and polypahryngeals:

Some pure populations were identified as mixed at the next collection (next years) and in the opposite season. However, **the existence of natural pure populations is uncertain – see the sampling remark (Material and method)**.

### 3.2 Morphology

#### The monopharyngeal *Crenobia alpine*

CR1, Cm1, Cr4, Cm2

These four specimens were collected in summer and are sexually matures, and, with very few exceptions, reveal ovaries, few prepharyngeal ventral follicular testes, ducts and fully developed copulatory apparatus. The exceptions are attributed to a physiological state or non-compliance with sectioning requirements (transversal sections in specimen Cm2 show ovaries, spermiducts, yet no copulatory apparatus because the worm was not entirely sectioned; the lack of ovaries and oviducts in specimen CR1 is considered a physiological state / a stage of the reproductive cycle).

##### The copulatory apparatus

Figs. 2, 3 (CR1), 4 – 8 (Cm1), 9 (CR4)

**Fig. 2.**
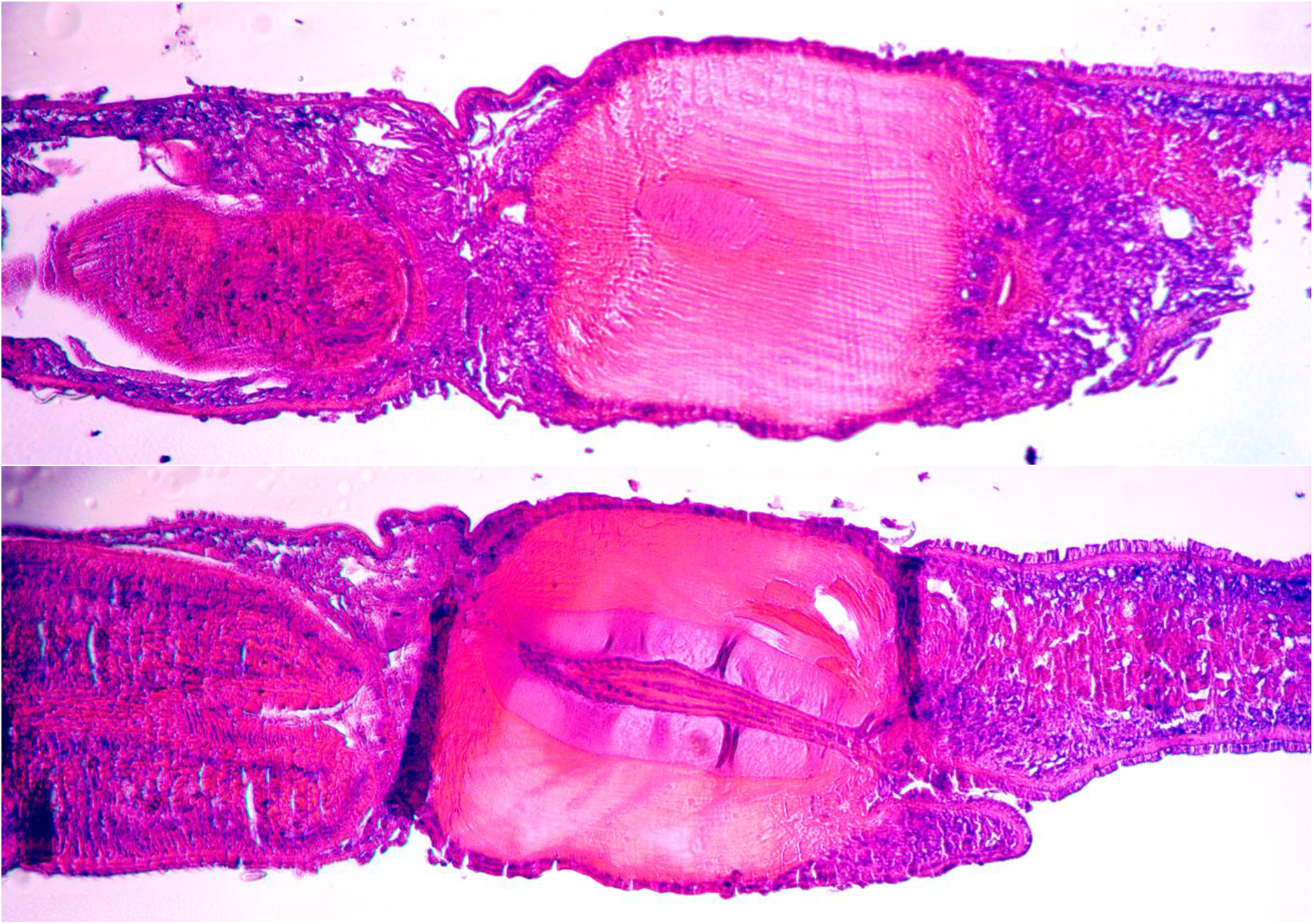
Specimen CR1, monopharyngeal, microphotographs of the copulatory apparatus

**Fig. 3.**
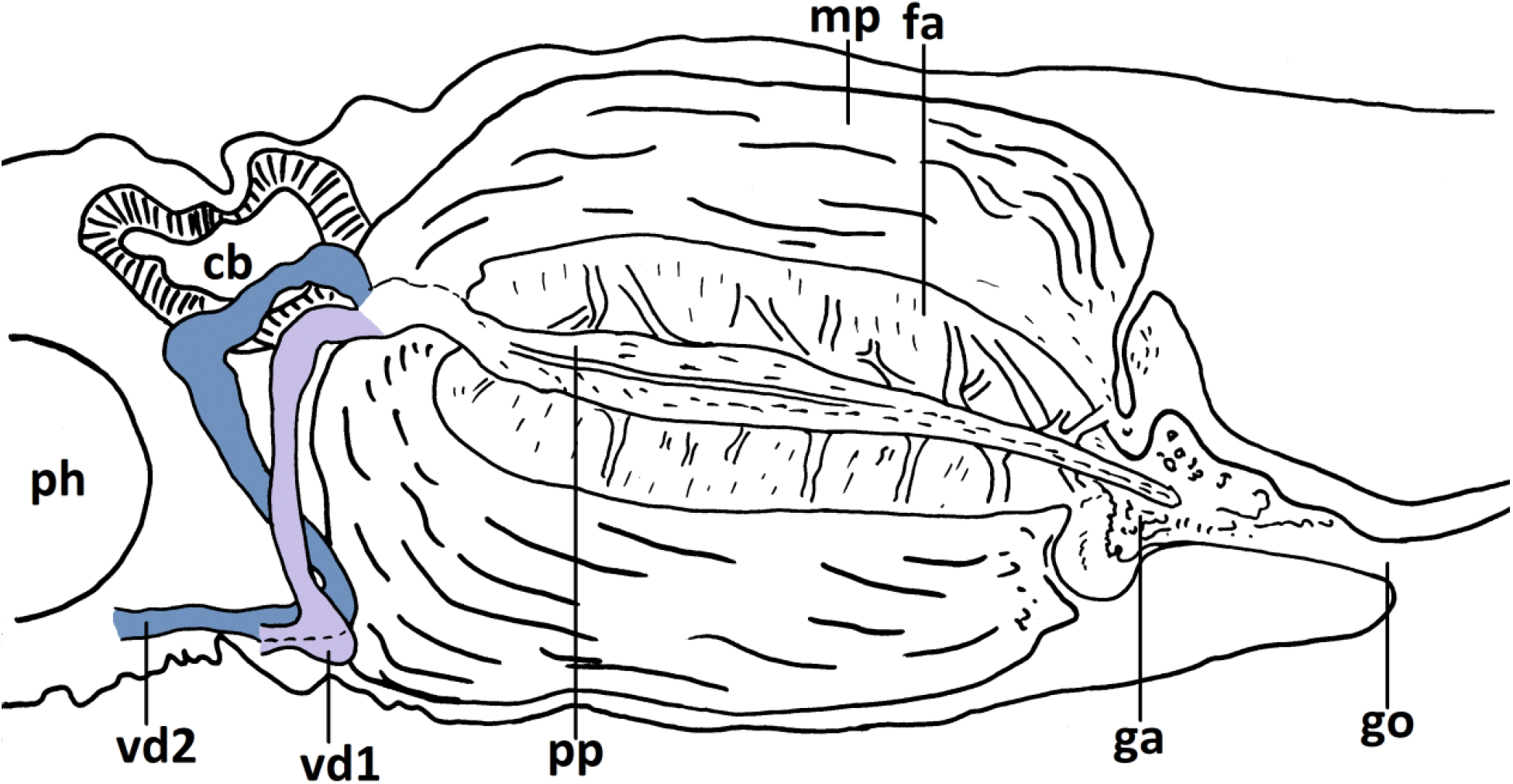
Specimen CR1, monopharyngeal, schematic reconstruction of the copulatory apparatus

**Fig. 4.**
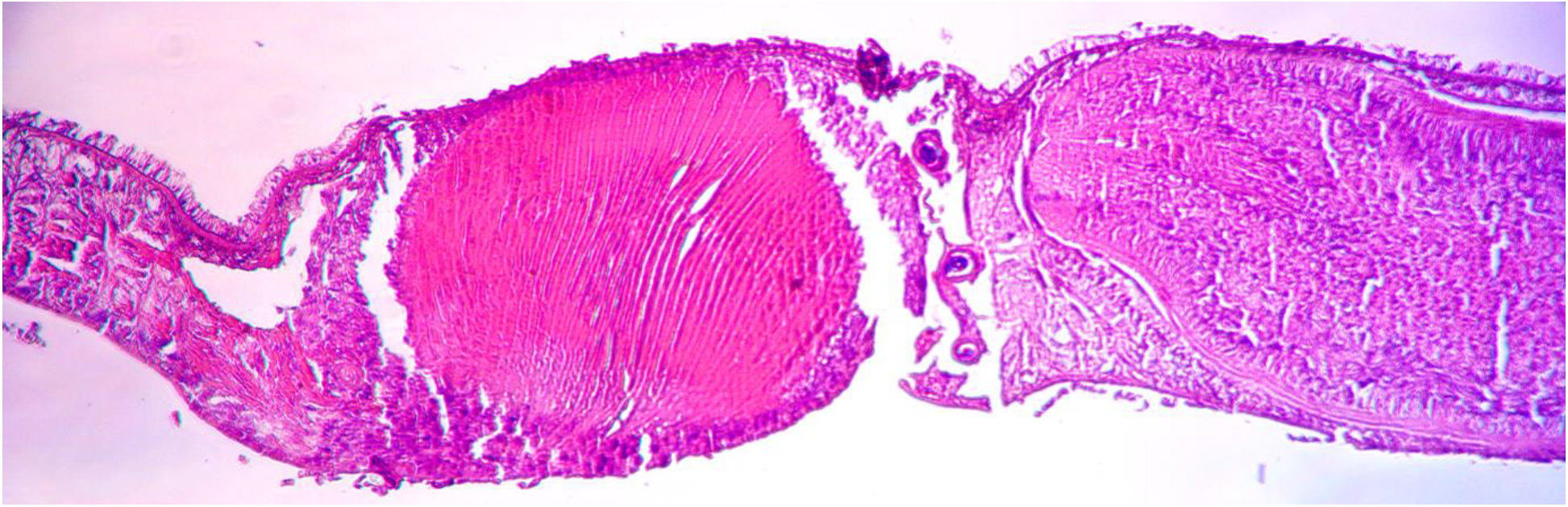
Specimen Cm1, monopharyngeal, microphotographs of vd1 course (proximal vasa deferens)

The copulatory apparatus follows the general pattern of the genus Crenobia:

- an overdeveloped (well developed) atrium consisting of 2 distinct parts: i) the external muscular part, very thick, composed by muscular plates (bundles) in radial disposition, surrounded by an outer zone of myoblasts (Sluys 2022) and ii) the internal fibrous layer
- the penis has an indistinct bulb and a long papilla
- the genital atrium is divided by an atrial fold in 2 regions: i) the proximal region which houses the most part of the penis papilla; it also receives the common oviduct and the bursal canal (Sluys 2022) and ii) a distal region which opens by the gonopore (the genital orifice).
- the two vasa deferentia (spermiducts) with large spermiducal vesicles in their terminal part, at a certain distance before opening into the penis bulb.

Besides the above general characters, the histological slides of the four analysed specimens reveal some particularities:

- the distal part of the genital atrium shows to be filled with a tissue (for instance Fig. 3)
- the course of the two vasa deferentia is symmetric in specimens CR1 and CR4 and asymmetric in specimen Cm1
- the presence of a complex system of apparently fine ducts or sclerotized lines in the wall of the genital atrium – Figs. 2, 5, 9. These structures seem to be arranged between, and interconnect the atrial space housing the penis papilla, the distal atrium, and the penis bulb. Correspondent orifices are visible in the penis bulb and the wall of the distal atrium.

**Fig. 5.**
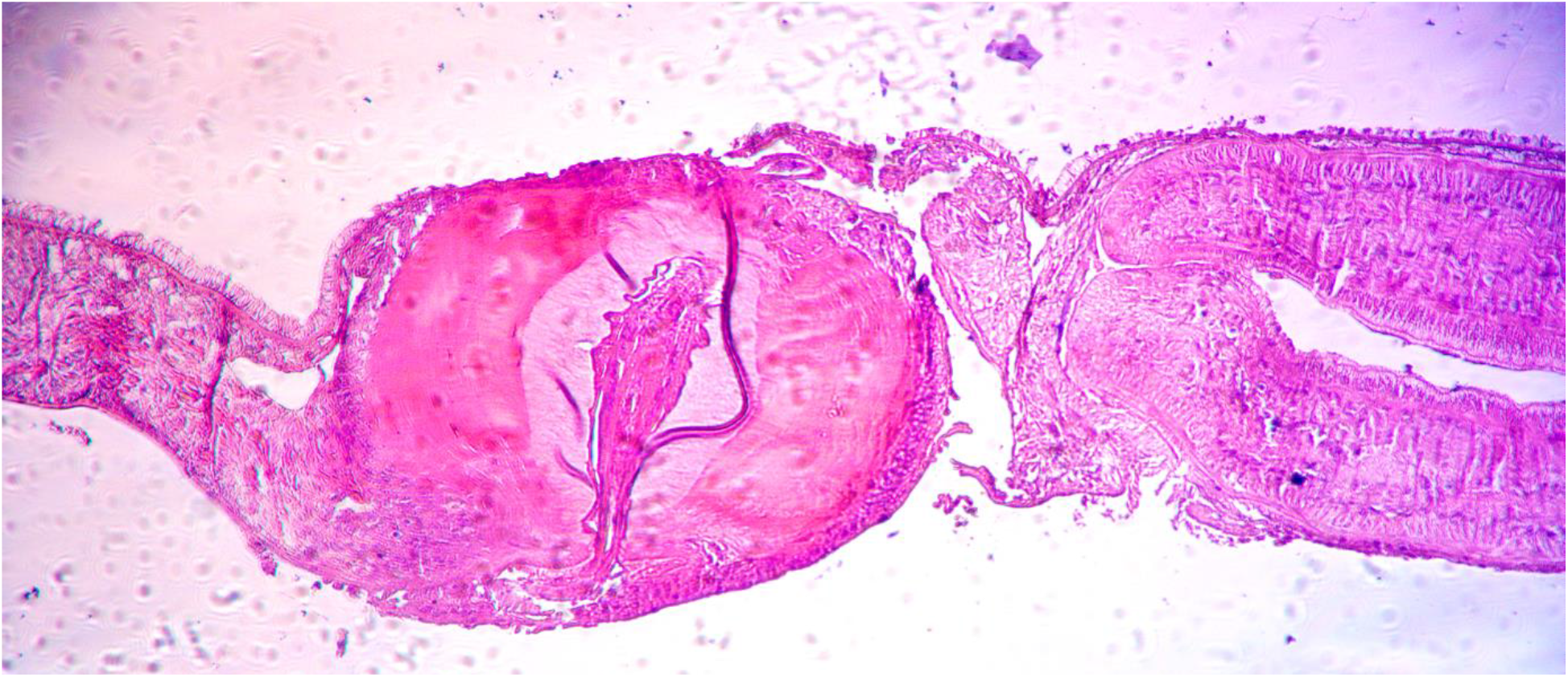
Specimen Cm1, microphotograph of the copulatory area with the penis papilla

**Fig. 6a.**
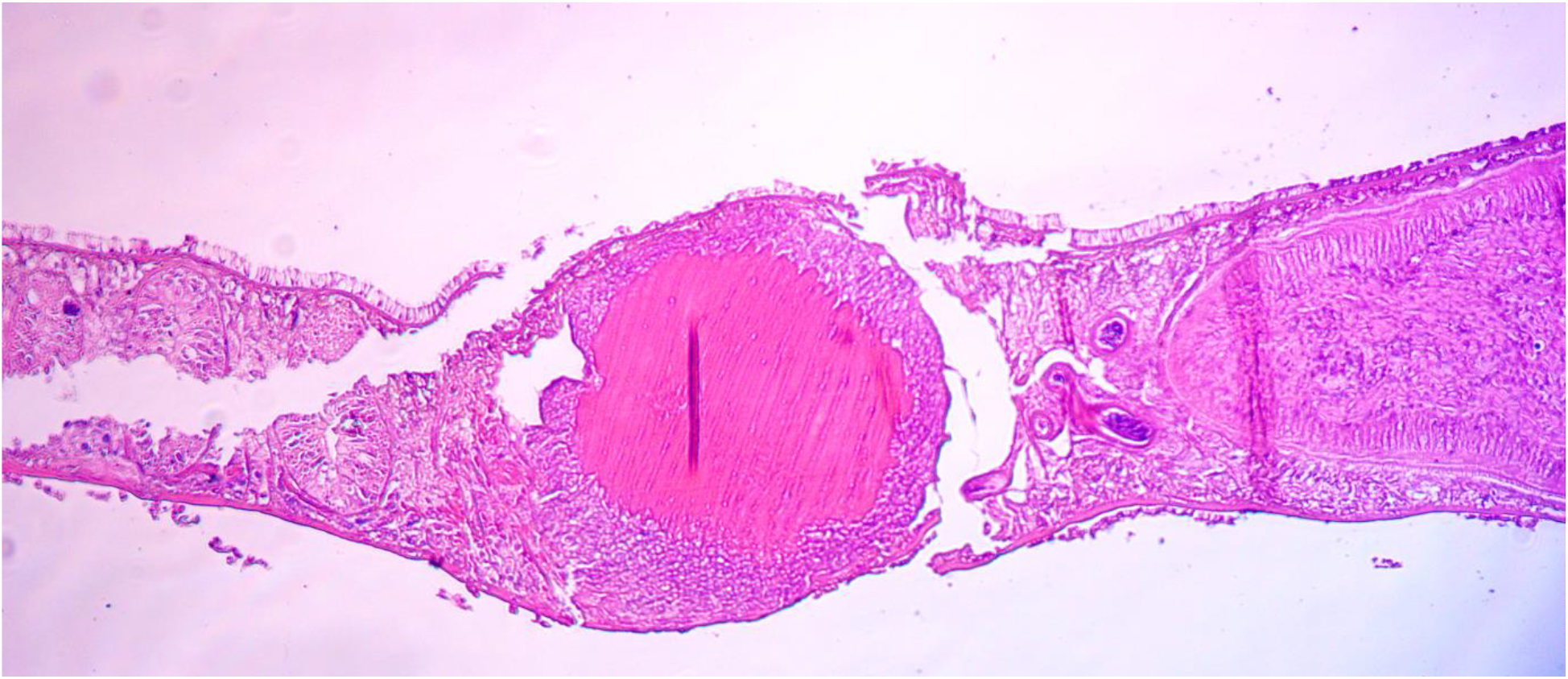
Specimen Cm1, microphotographs of vd2 course (distal vasa deferens)

**Fig. 6b.**
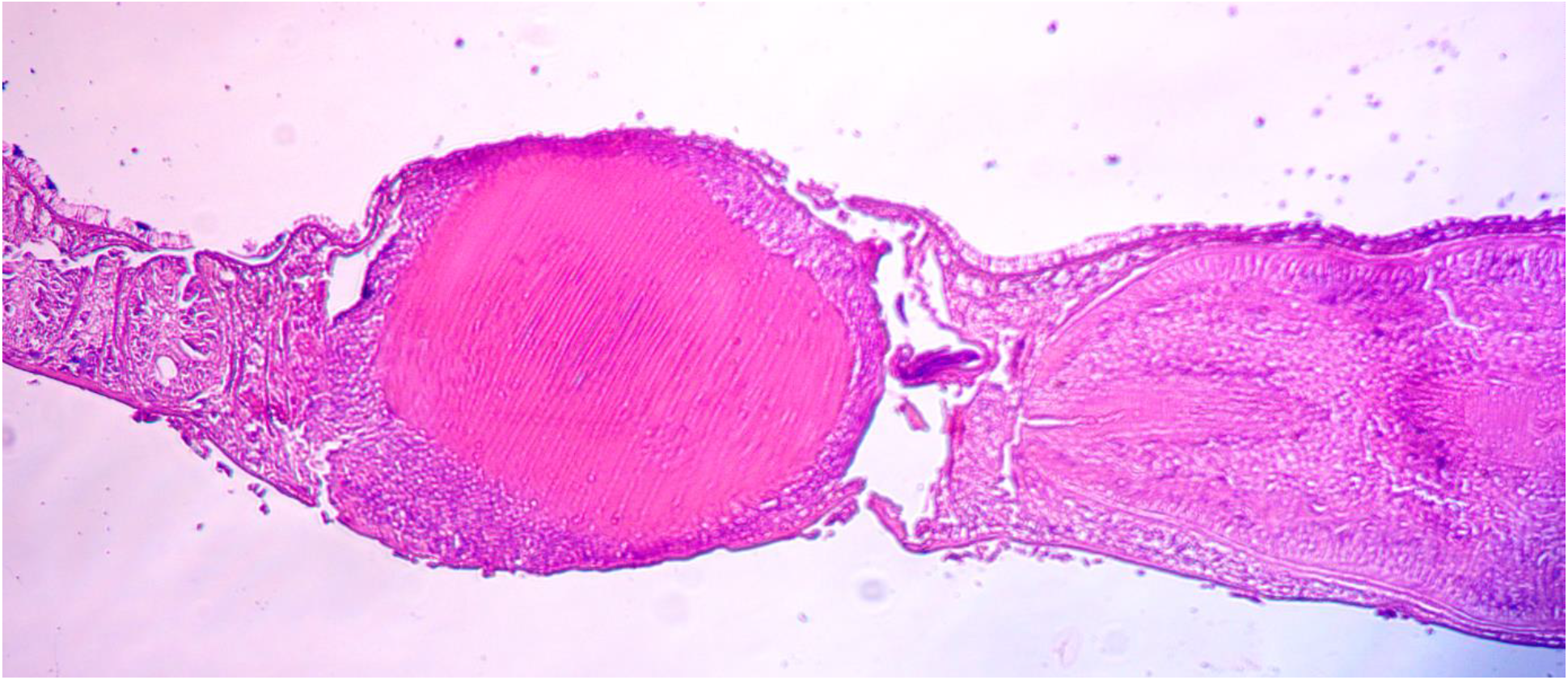
Specimen Cm1, microphotographs of vd2 course (distal vasa deferens)

**Fig. 7.**
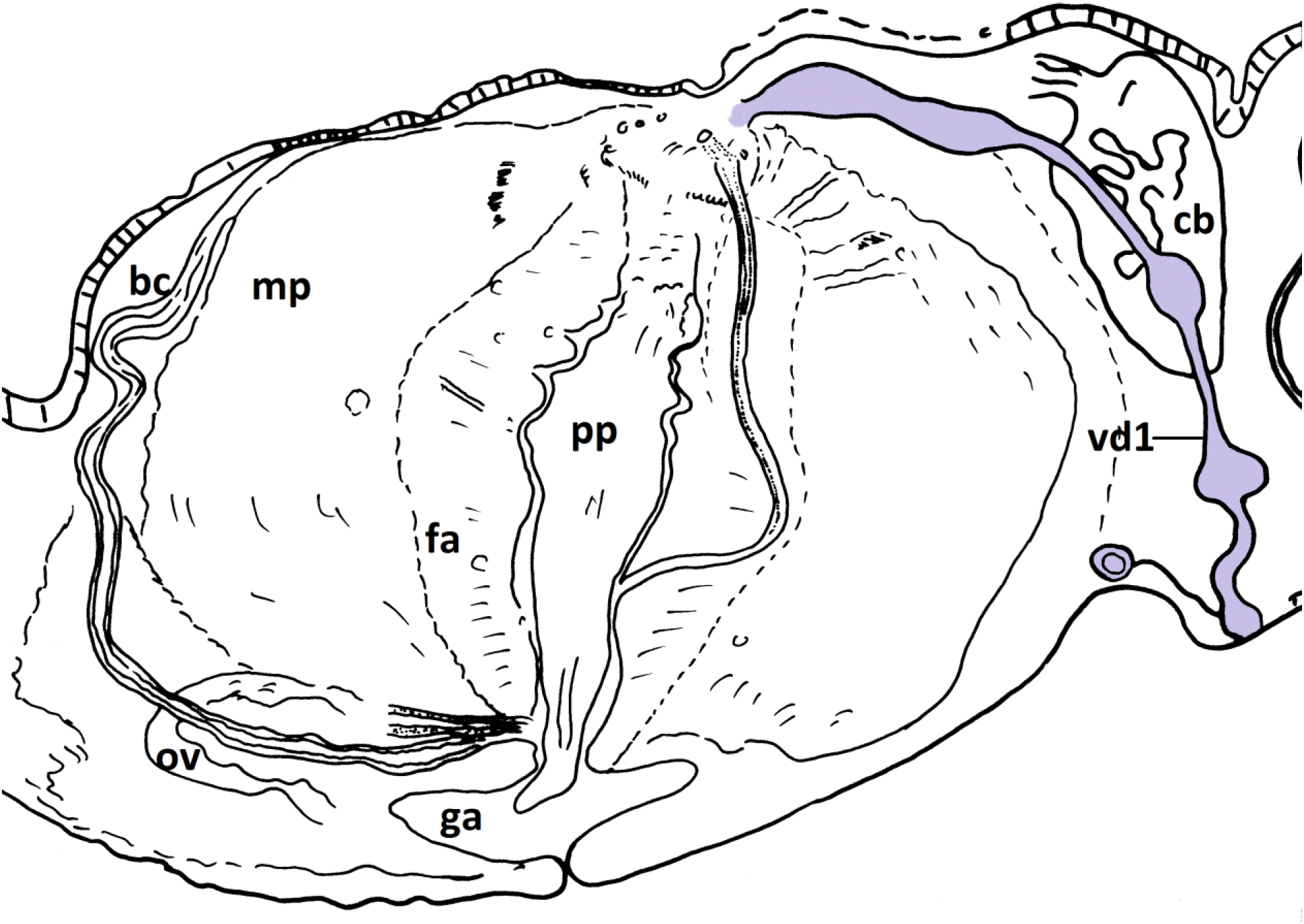
Specimen Cm1, schematic reconstruction of the copulatory apparatus with the course of vd1 (proximal vasa deferens)

**Fig. 8.**
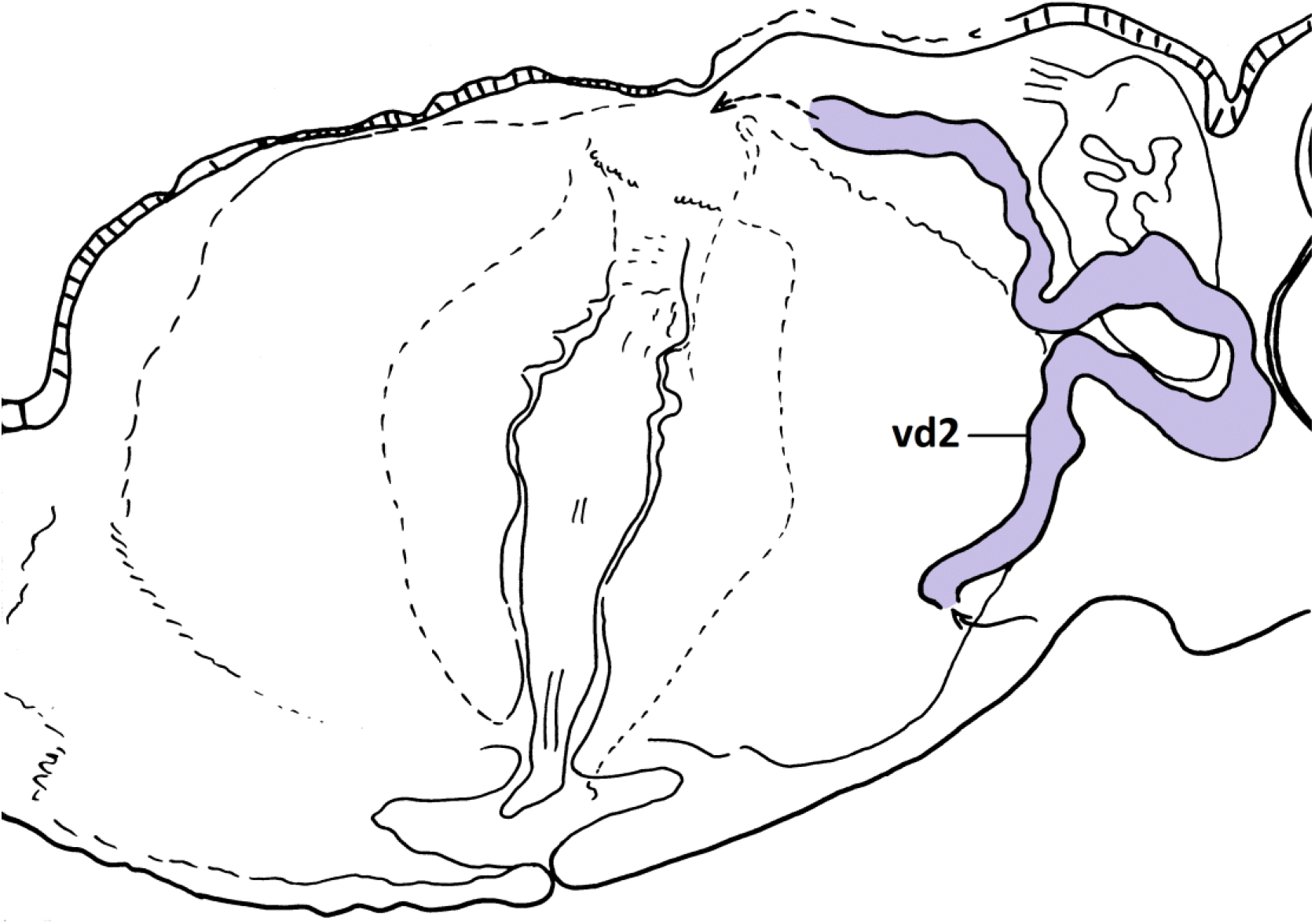
Specimen Cm1, schematic reconstruction of the copulatory apparatus with the course of vd2 (distal vasa deferens)

**Fig. 9.**
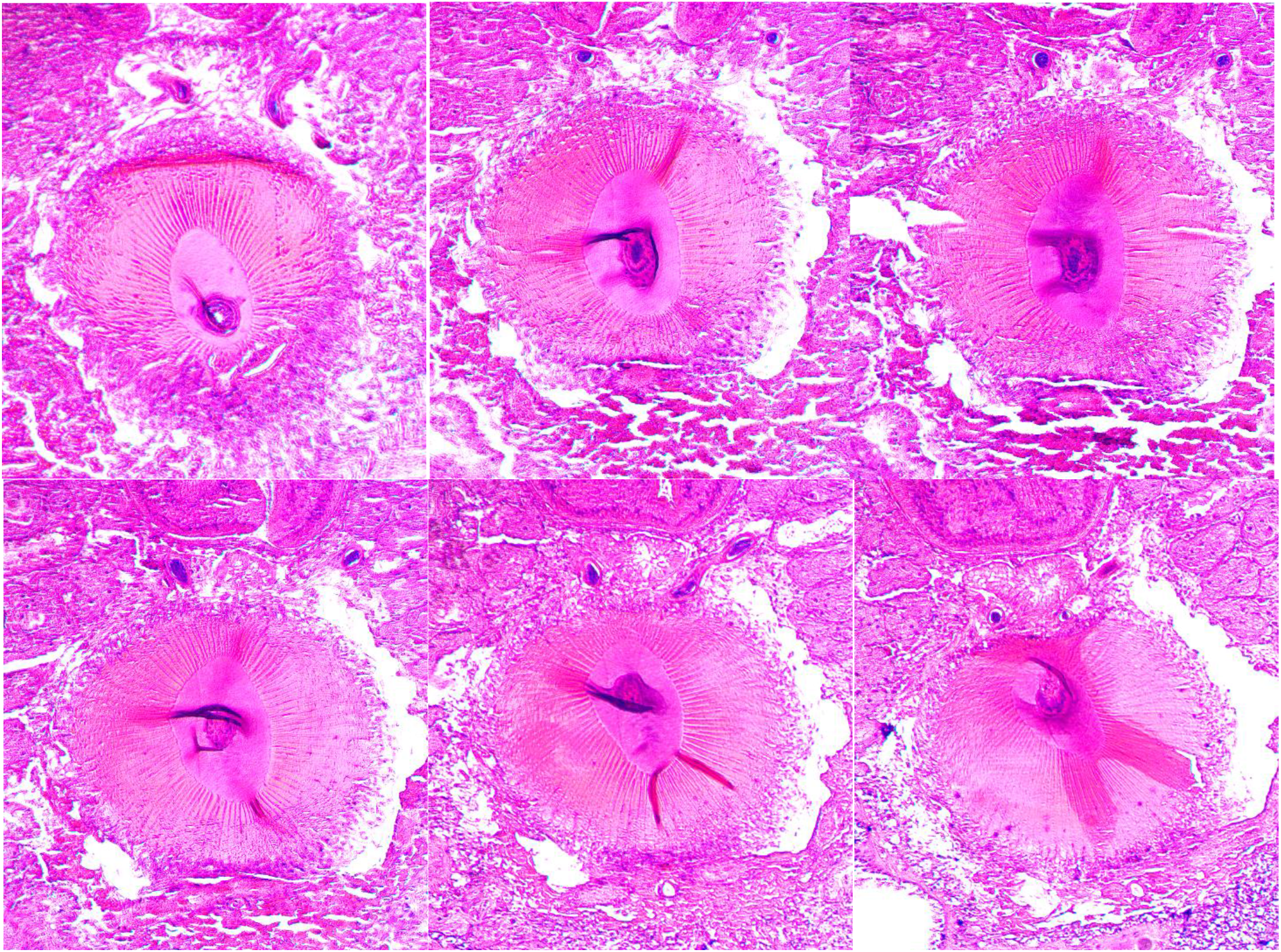
Specimen CR4, monopharyngeal, frontal section under an angle, aspects of the copulatory apparatus with the system of fine ducts/sclerotized lines

#### The polypharyngeal *Crenobia montenigrina*

(CR8 – sexually mature; CR2, CR3, Cp1, Cp2 - immatures)

Besides the pigmented specimens, unpigmented (white) and marbled polypharyngeals were also collected – Fig. 1 Appendix

##### The pharynxes

– numerous; every pharynx has its own root and opening; the pharynxes open into a common space which has one posterior mouth at a certain distance from the genital orifice) Fig. 10 (CR8-37.2, CR2-24.1).

**Fig. 10.**
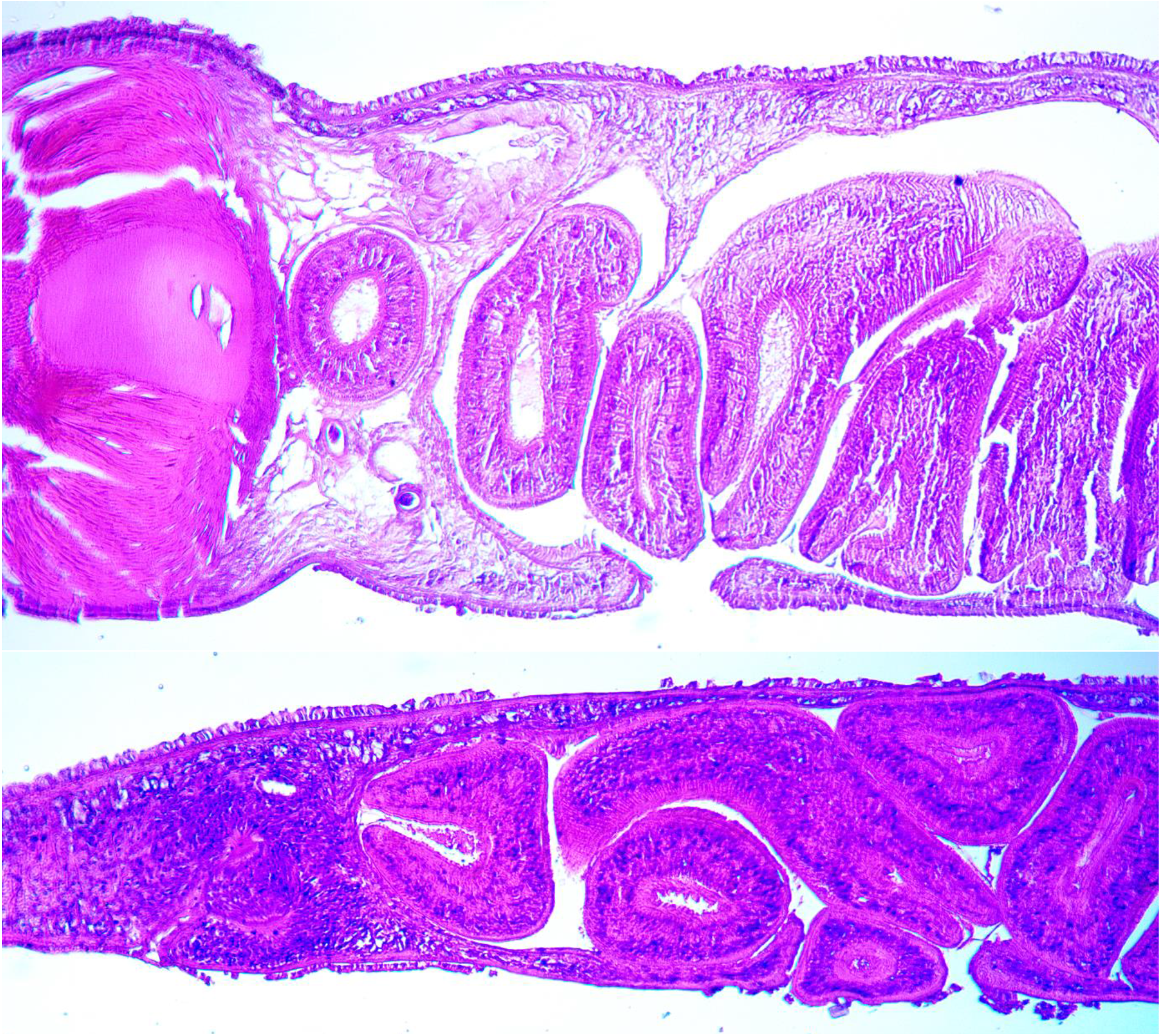
Mouth opening: up – specimen CR8, polypharyngeal, mature, mouth (slide CR8-37-2); down – specimen CR2 (slide.24.1), polypharyngeal, immature – mouth (right) and gonopore (left)

##### The testes and ovaries

ovaries present in sexual CR8 (CR8-37.1); small and ventral rare follicular testes, most of them located prepharyngeal (CR8-24.1, CR8-29.1, CR8-37.3), only one case located interpharyngian and anterior (between the anterior pharynxes – CR8-29.1). The immature CR3 reveals: i) intense basophilic clusters of cells, possibly immature follicular testes located anterior and lateral to the first pharynx – Fig. 11. The ante-pharyngian cluster of possible follicular testes are surrounded by even more basophilic mesenchymal cells and ii) a cluster of cells, possibly developing ovary located in the cephalic region - Fig. 12. The simultaneous development of testes and ovaries indicates neither proterandry, nor protogyny. No ovaries, nor testes observed in specimens Cp2.

**Fig. 11.**
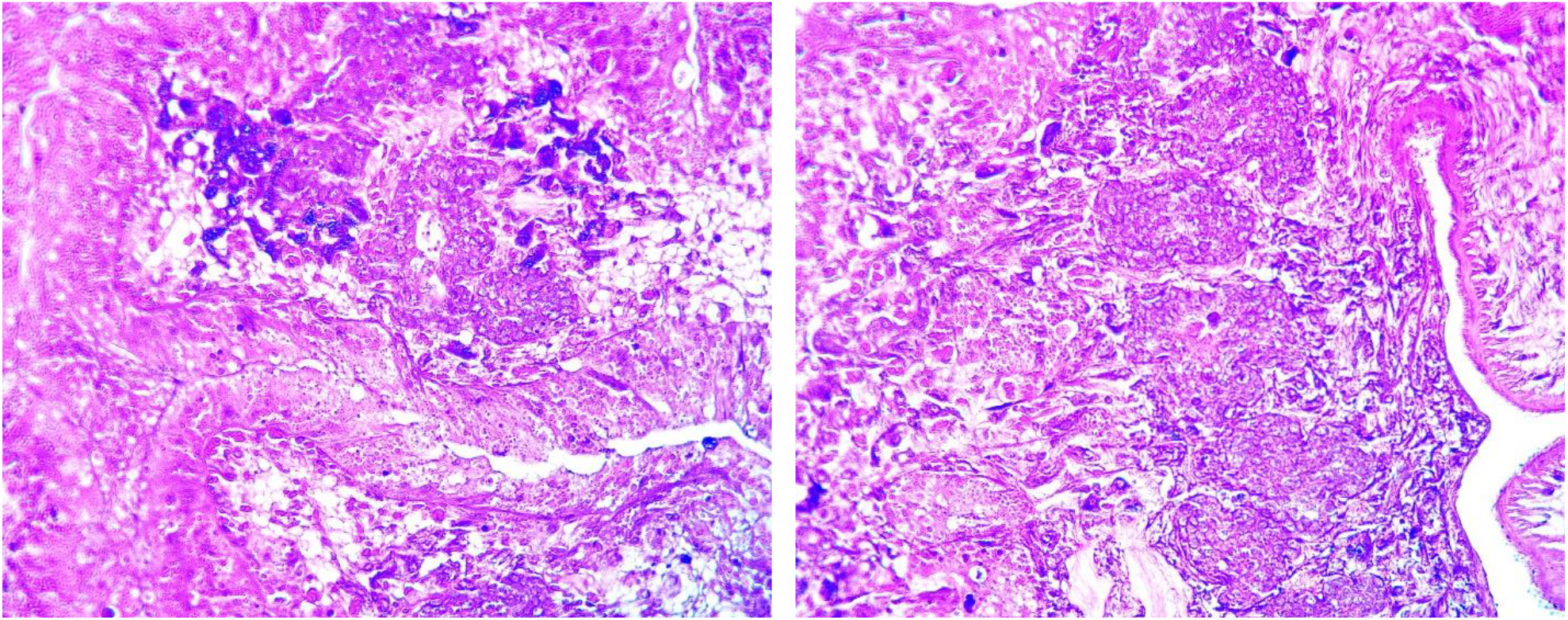
Specimen CR3 – immature, polypharyngeal, developing follicular testes? in prepharyngian position (left image) and lateral to the pharynx (right image) (slide CR3-3-2)

**Fig. 12.**
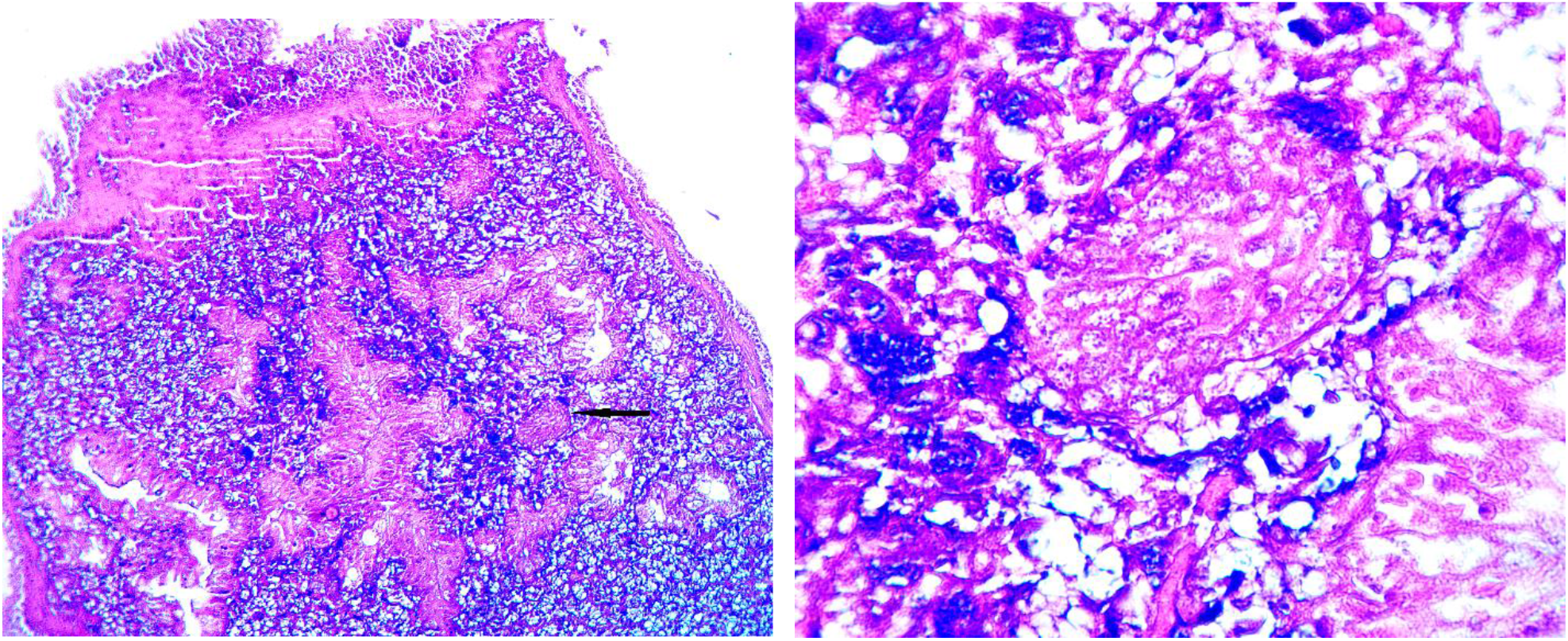
Specimen CR3 – immature, polypharyngeal, developing ovary? (slide CR3-6-1)

##### The copulatory apparatus

Shows to be incompletely developed in specimens collected in summer: Cp1, Cp2 (slide Cp2-55.4), CR3 (slide CR3-4.2) and CR2 (slide CR2-24.1) – Fig. 13.

**Fig. 13.**
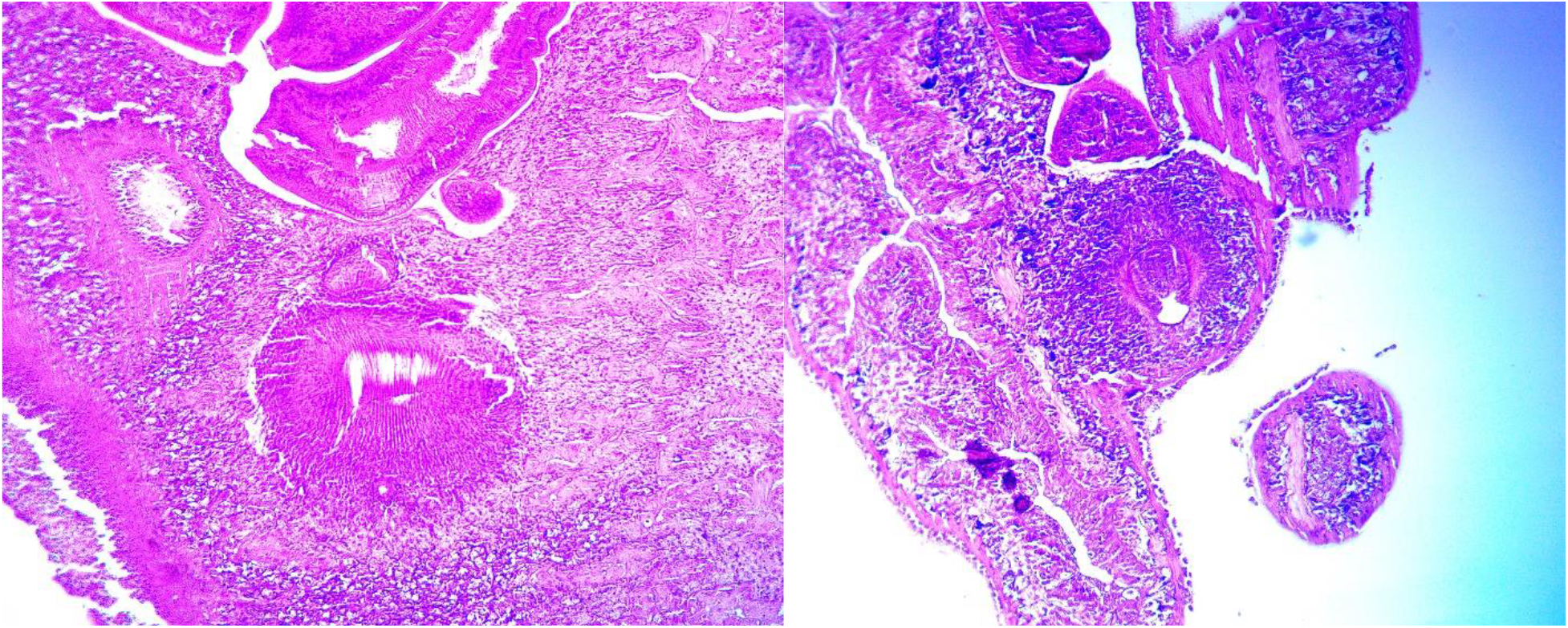
Immature copulatory area in polypharyngeals Left: specimen Cp2 – (slide Cp2-5-4); right: specimen CR3 (slide CR3-4-2)

For the sexual CR8 collected in November, besides the general characters, the histological slides reveal the following particularities – Figs. 14 - 18:

**Fig. 14a.**
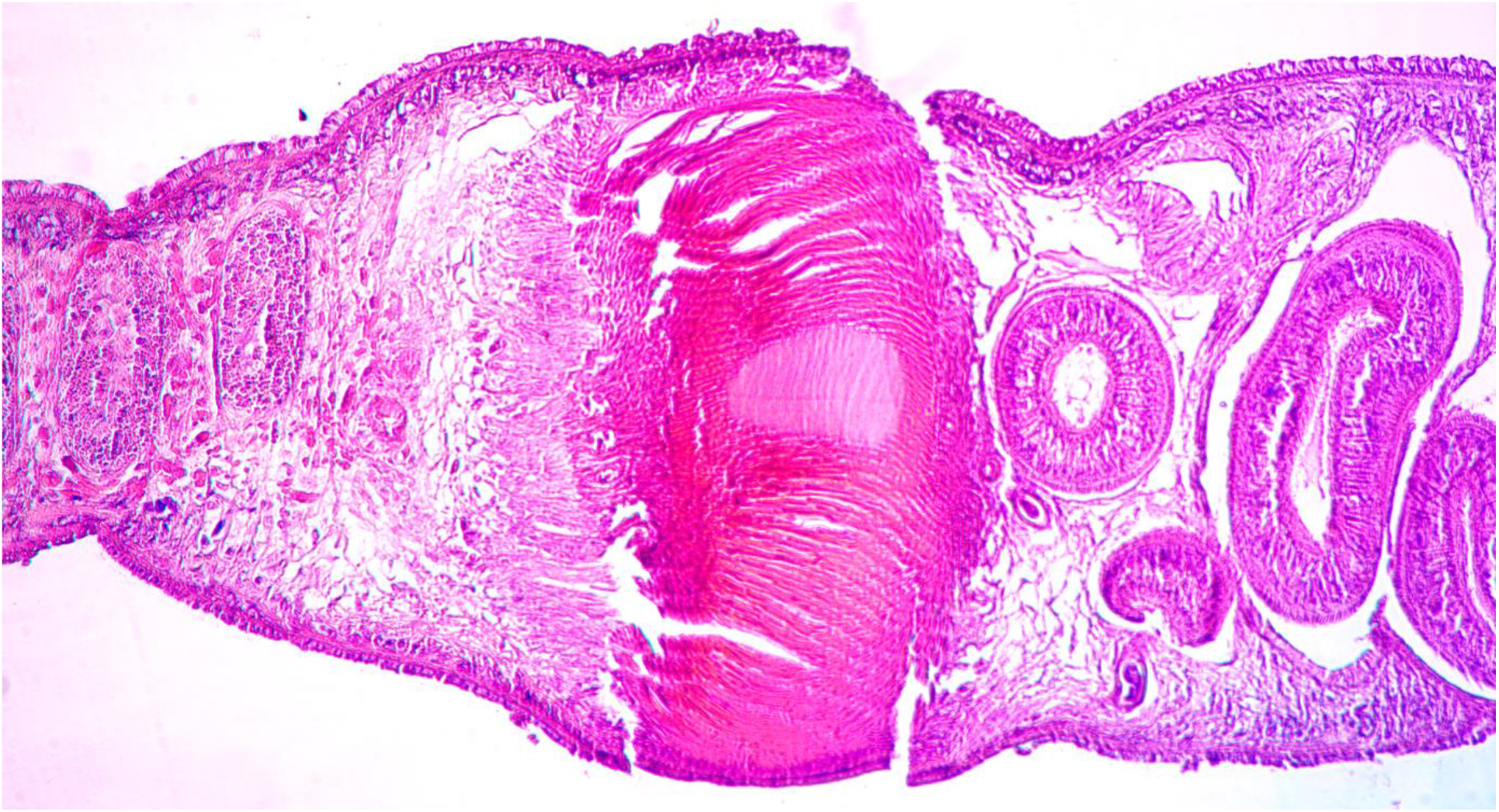
Specimen CR8, polypharyngeal, microphotographs of the copulatory apparatus with vd1 course (proximal vasa deferens)

**Fig. 14b.**
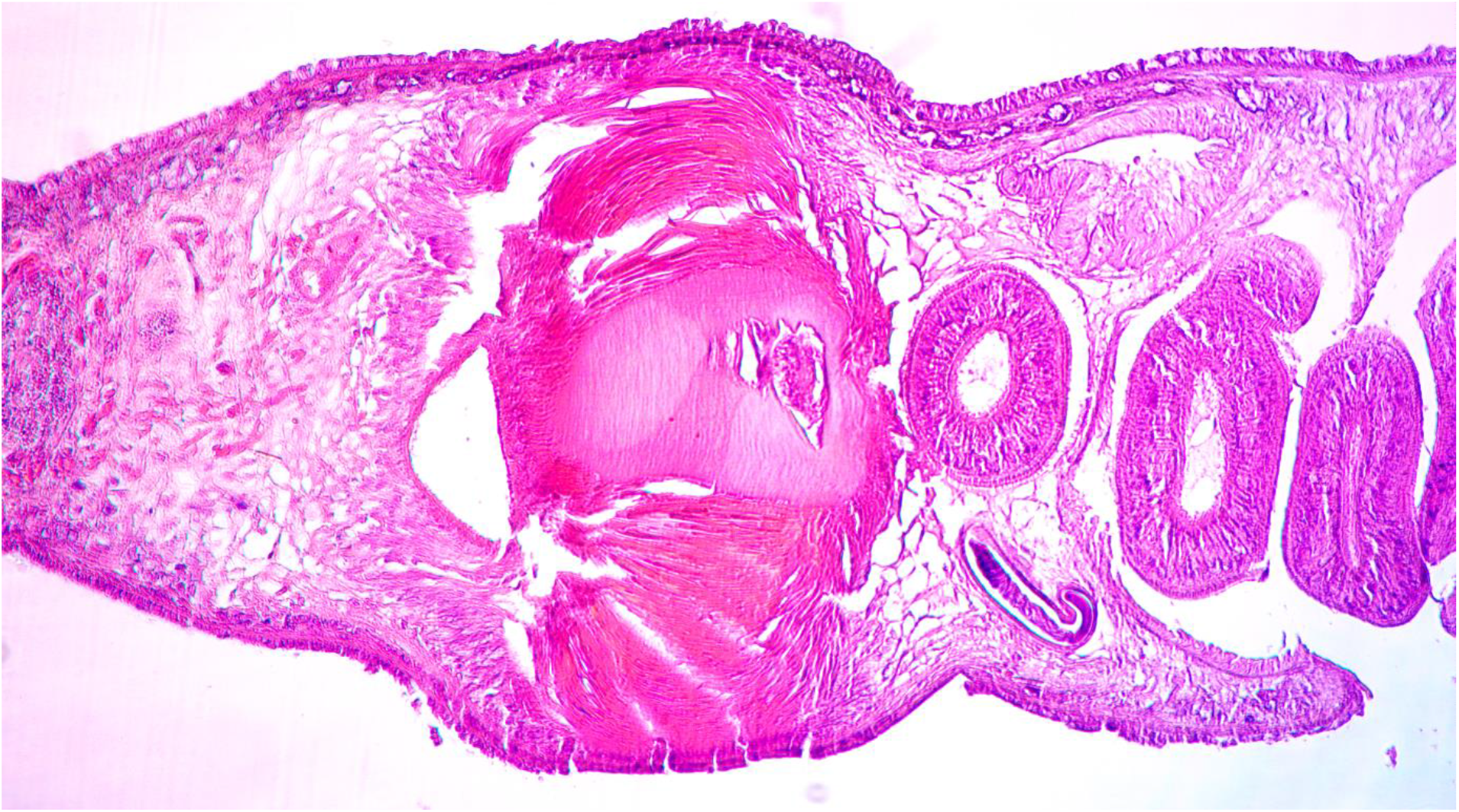
Specimen CR8, polypharyngeal, microphotographs of the copulatory apparatus with vd1 course (proximal vasa deferens)

- the distal part of the genital atrium filled with a tissue – Fig. 15

**Fig. 15.**
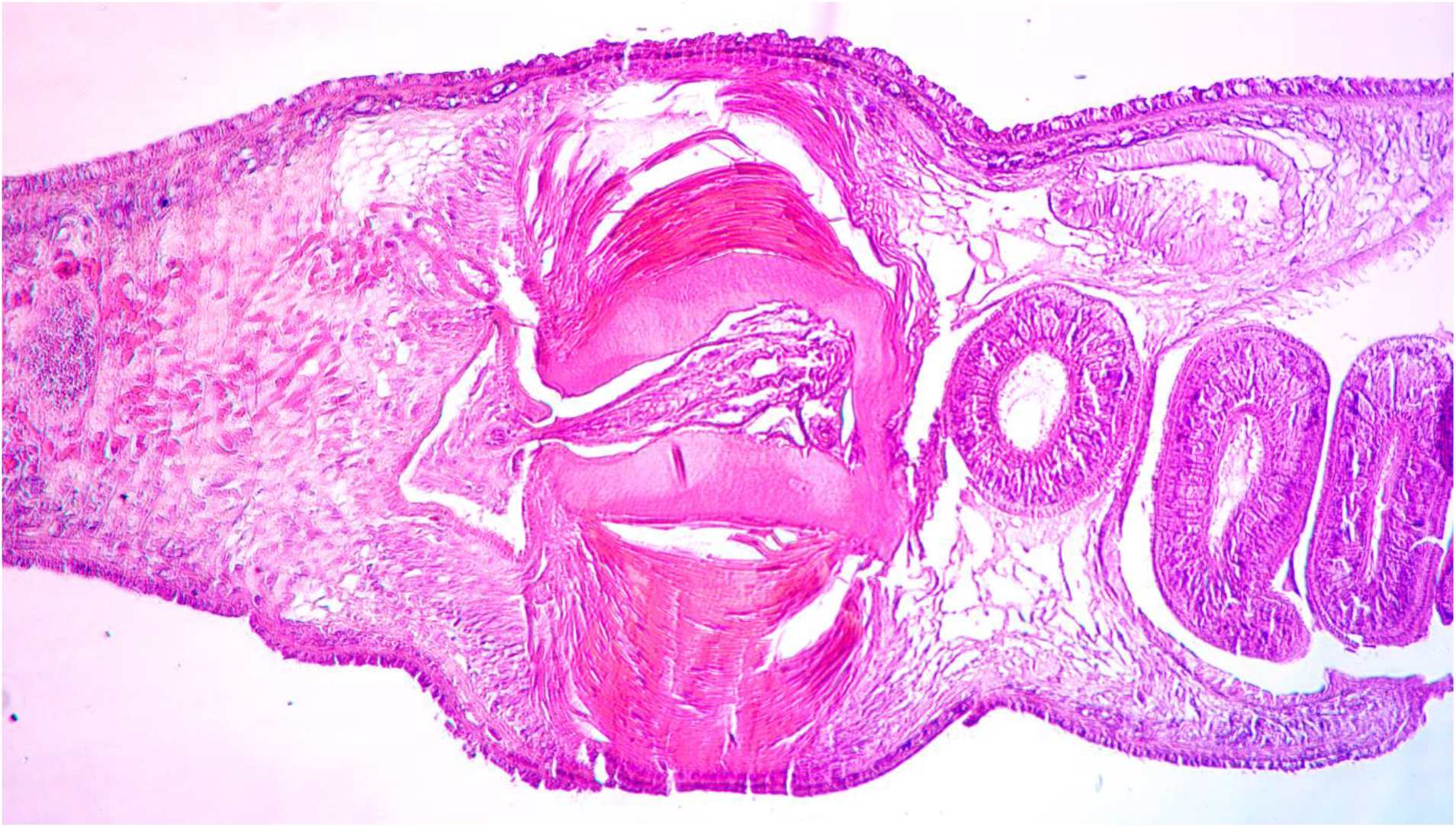
Specimen CR8, polypharyngeal, microphotograph of the copulatory apparatus with the penis papilla

**Fig. 16a.**
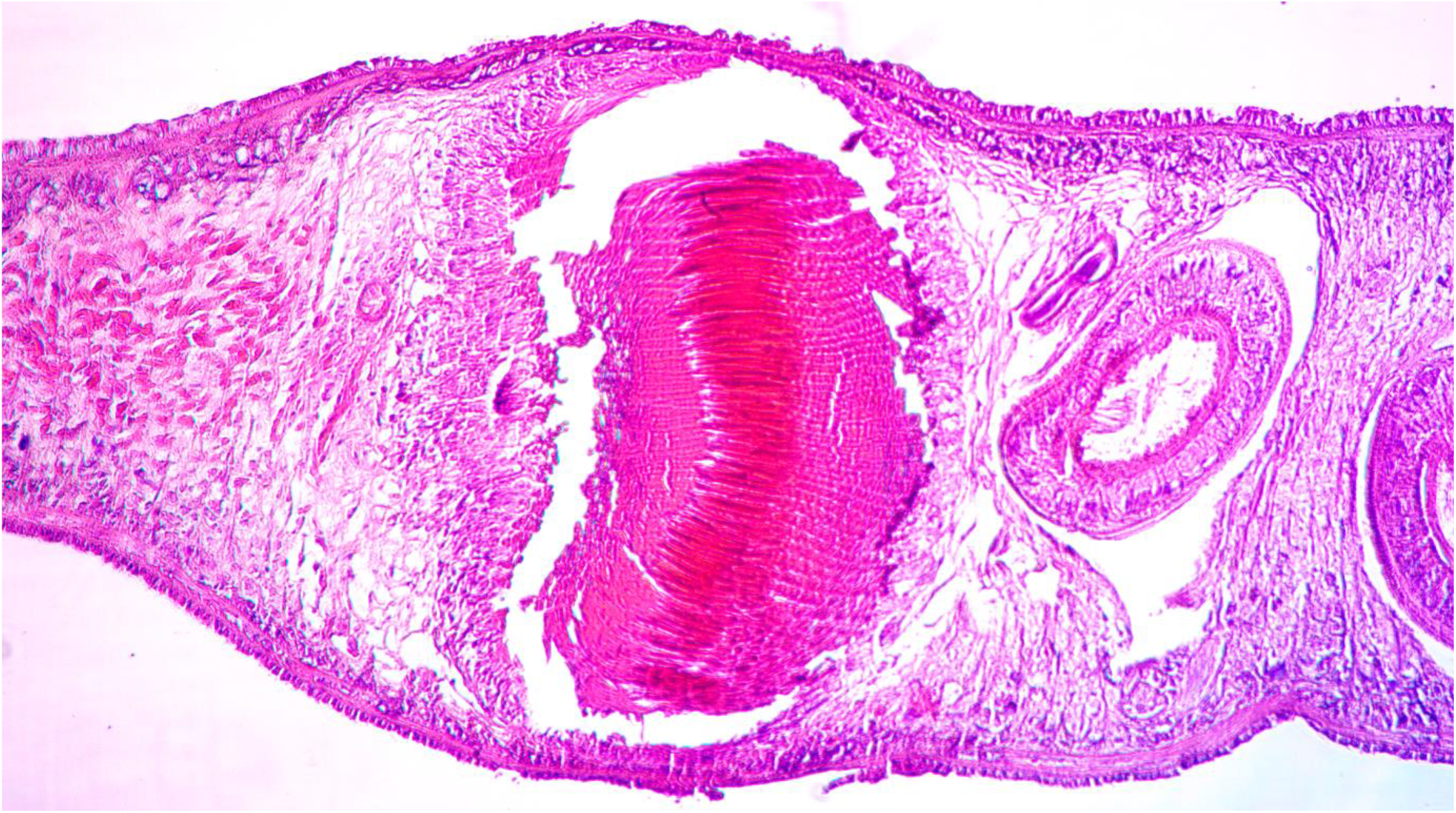
Specimen CR8, polypharyngeal, microphotograph of the copulatory apparatus with vd2 course (distal vasa deferens)

**Fig. 16b.**
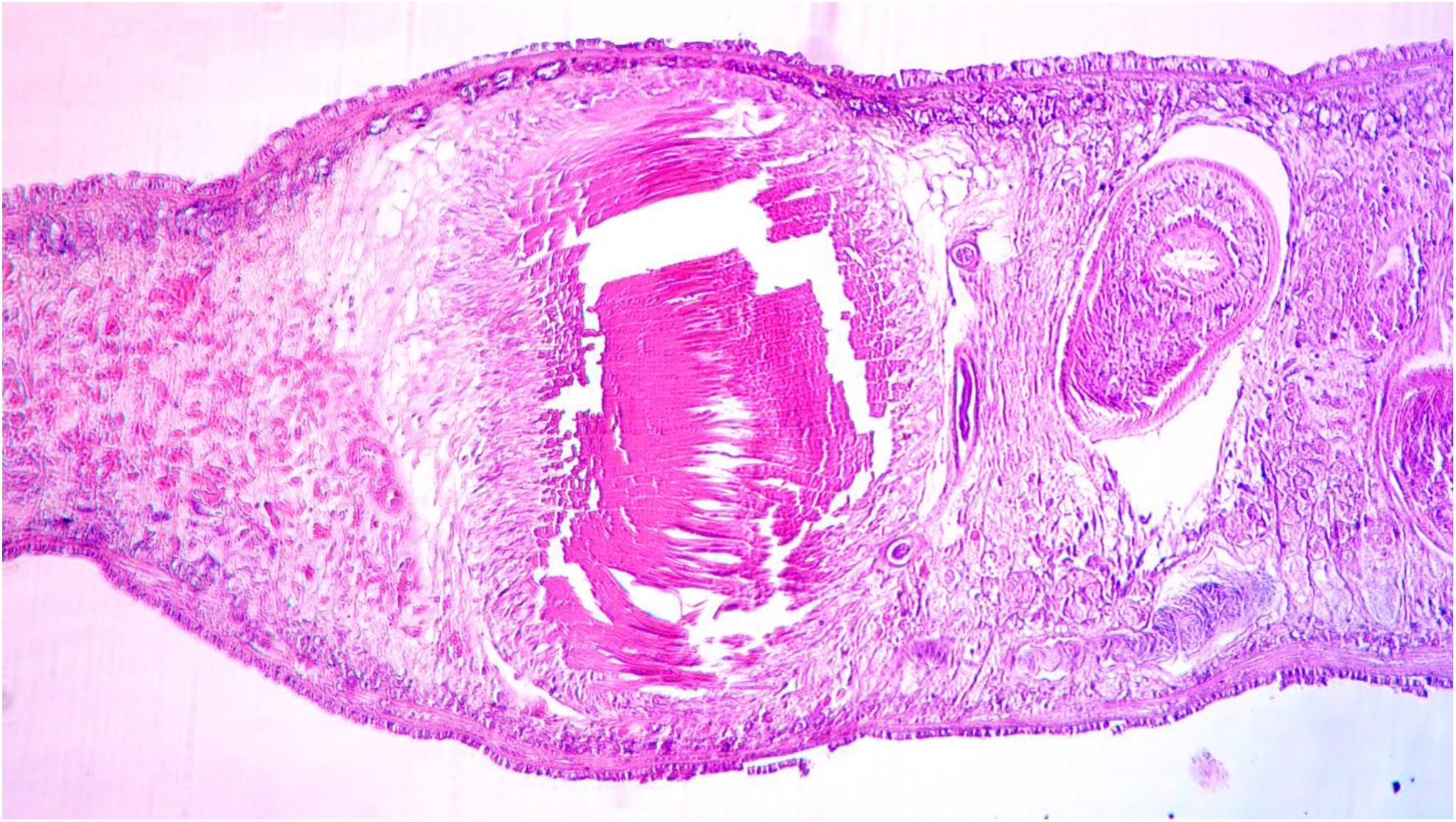
Specimen CR8, polypharyngeal, microphotograph of the copulatory apparatus with vd2 course (distal vasa deferens)

**Fig. 17.**
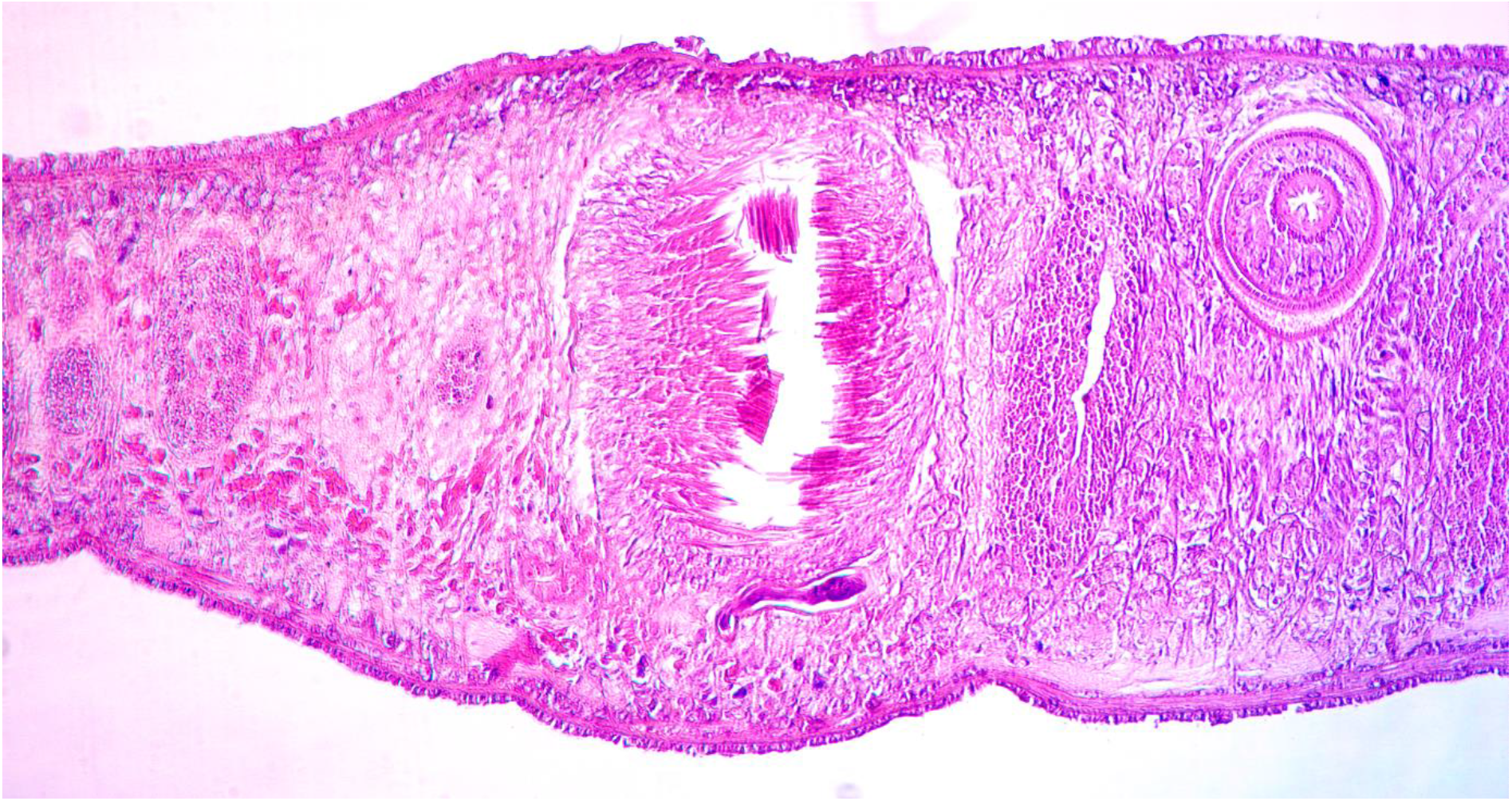
Specimen CR8, polypharyngeal, microphotograph of the copulatory apparatus with vd2 course (distal vasa deferens)

**Fig. 18.**
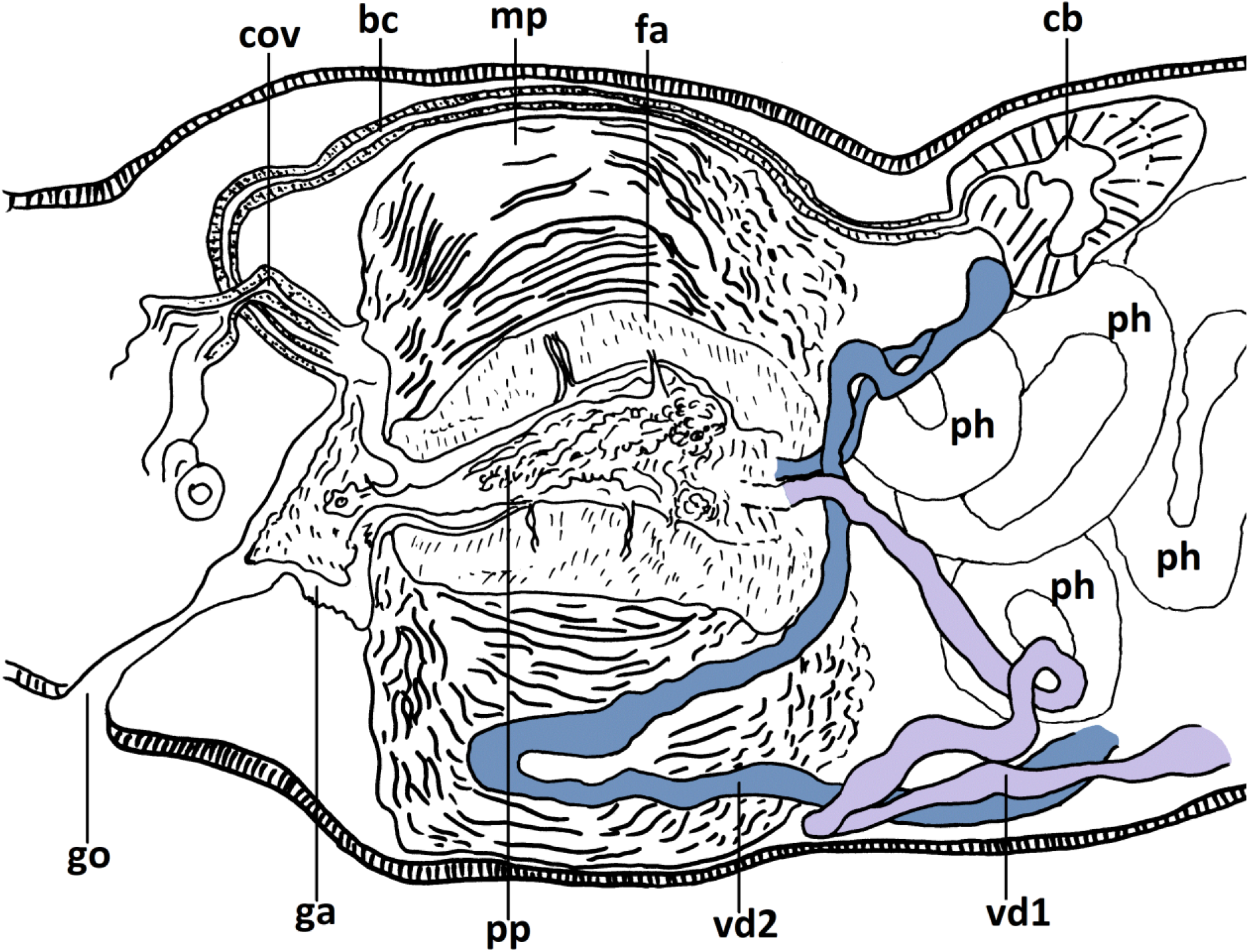
Specimen CR8, polypharyngeal, schematic reconstruction of the copulatory apparatus

- the asymmetry of the vasa deferentia
- the system of fine ducts is present but less developed compared with the monopharyngeals

### 3.3 Aspects regarding the reproductive biology (Figures 19 – 22 and Table 1)

For both forms, the analysis was focused on mixed populations.

**Table 1:** Comparative synthetic table of the copulatory area maturity of the monopharyngeals and polypharyngeals in mixed populations during the warm and cold seasons.

|  | Copulatory area<br>Warm season | Copulatory area<br>Cold season |
| --- | --- | --- |
| Monopharyngeal specimens | - large copulatory area in most specimens<br>- internal cocoons (copulatory areas with the aspect of internal cocoon) | - small copulatory area and<br>- asexuals (copulatory area absent) |
| Polypharyngeal specimens | - small copulatory area<br>- absent copulatory area<br>- undergo fission | - large copulatory area<br>- no cocoons observed<br>- undergo fission |

**Fig. 19.**
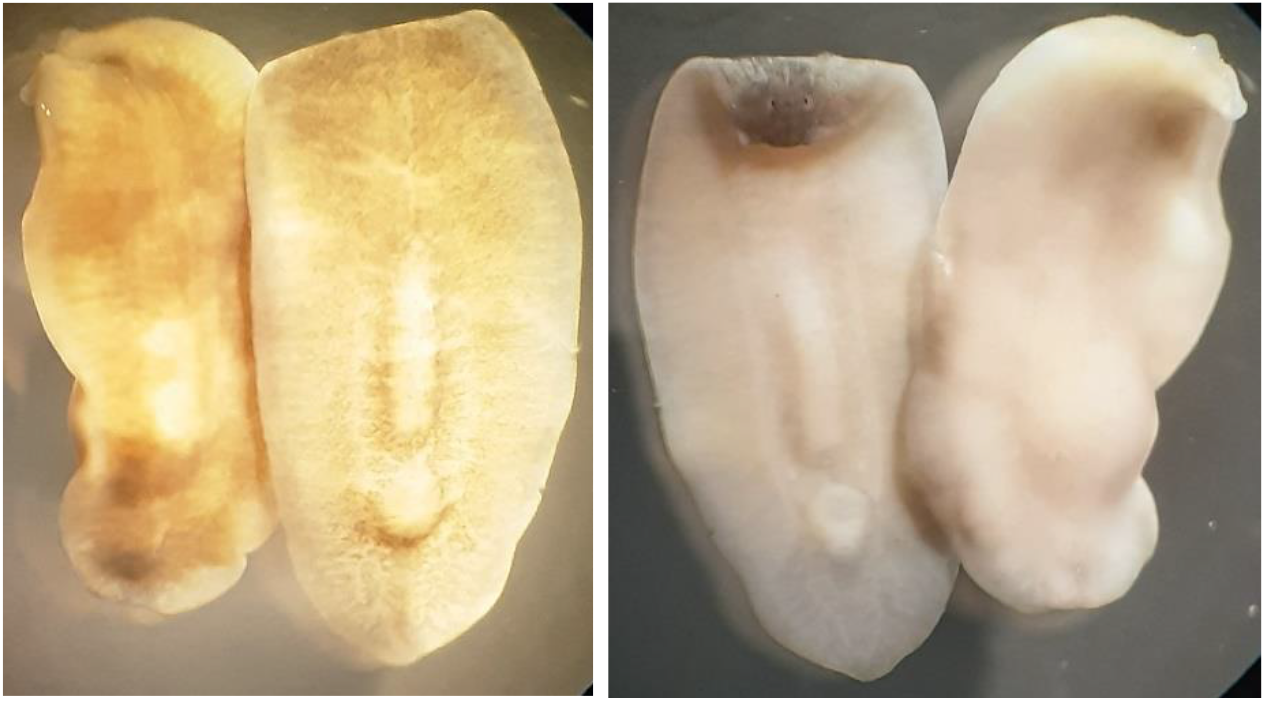
Monopharyngeal specimen – 5 iul. 2021, copulatory area with cocoon appearance (internal cocoon)

#### The monopharyngeal *Crenobia alpina*

The analysis of the fixed uncut specimens and of the histological slides shows aspects of the reproductive biology:

##### A) During the warm season (June, July)

- most part of the monopharyngeals have large copulatory area, fully developed for sperm transfer during mating (CR1, CR4, Cm1 and visual inspection of uncut specimens). One specimen presented an overdeveloped copulatory area with the aspect of an internal cocoon – Fig. 19. No external cocoons have been observed. An extremely interesting observation was done in the sample of the reocrene no. 3 prepared for 96º ethanol (starvation of the specimens for 5 days). The individuals of this sample were kept for 5 days in the container in which they were collected. At the fixation, 4 – 5 newly hatched specimens were observed but unfortunately, they were lost (by negligence).
- very few specimens with asexual appearance
- few specimens with small copulatory area

Also, from pure monopharyngeal populations (teleski area) one specimen with cocoon aspect of the copulatory area was collected – Fig. 20 (left).

**Fig. 20.**
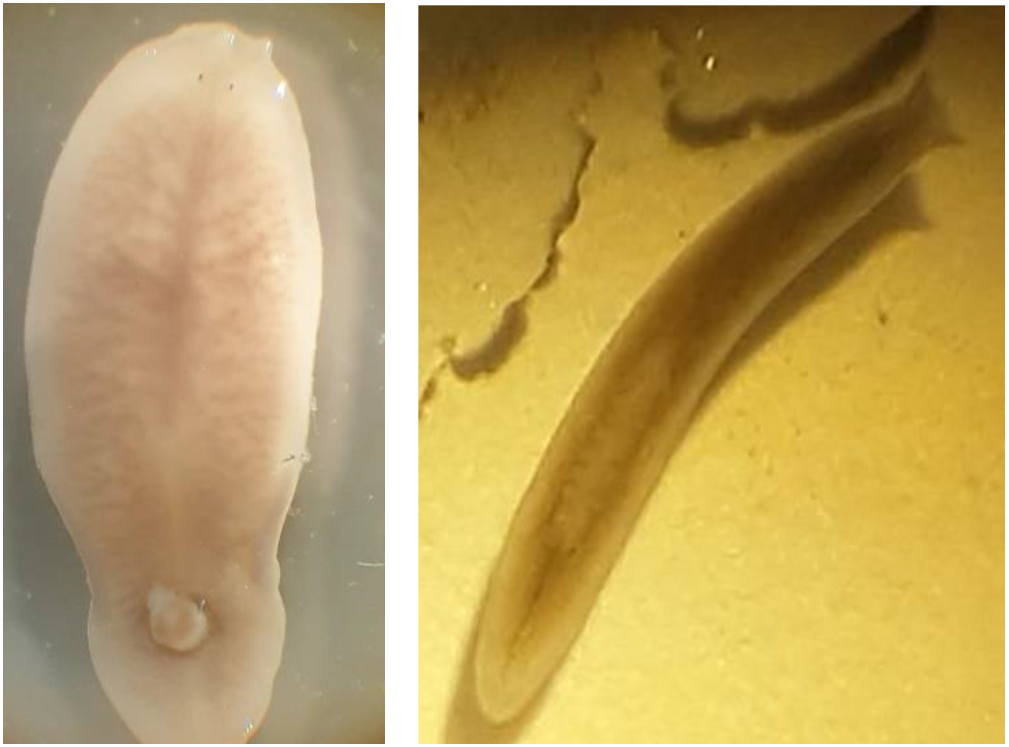
Monopharyngeals – left: 28.06.2017 (teleski area), internal cocoon exposed on the dorsal surface; right: asexual specimen, very small size (7.11.2022, reocren 2), most probably resulted from hatching, note the round posterior end.

##### B) During the cold season (November)

- part of the monopharyngeals seem to be uncapable for sperm transfer, they have small copulatory area (underdeveloped)
- part of the monopharyngeals are asexual, they lack the copulatory area – Fig. 20 (right).

#### The polypharyngeal *Crenobia montenigrina*

The reproductive state of the polypharyngeals in mixed populations:

A. During the warm season (June, July) most specimens have small copulatory area, and some are asexual (they lack the copulatory area).
B. During the cold season (November)
  - all specimens with large copulatory area (reocren 3, 7.11.2022); all asexuals (no copulatory area) (reocren 2, 7.11.2022)

The polypharyngeals were found undergoing fission during July and November in pure populations – Fig. 21, and specimens resulted from fission were sampled – Fig. 22.

**Fig.21.**
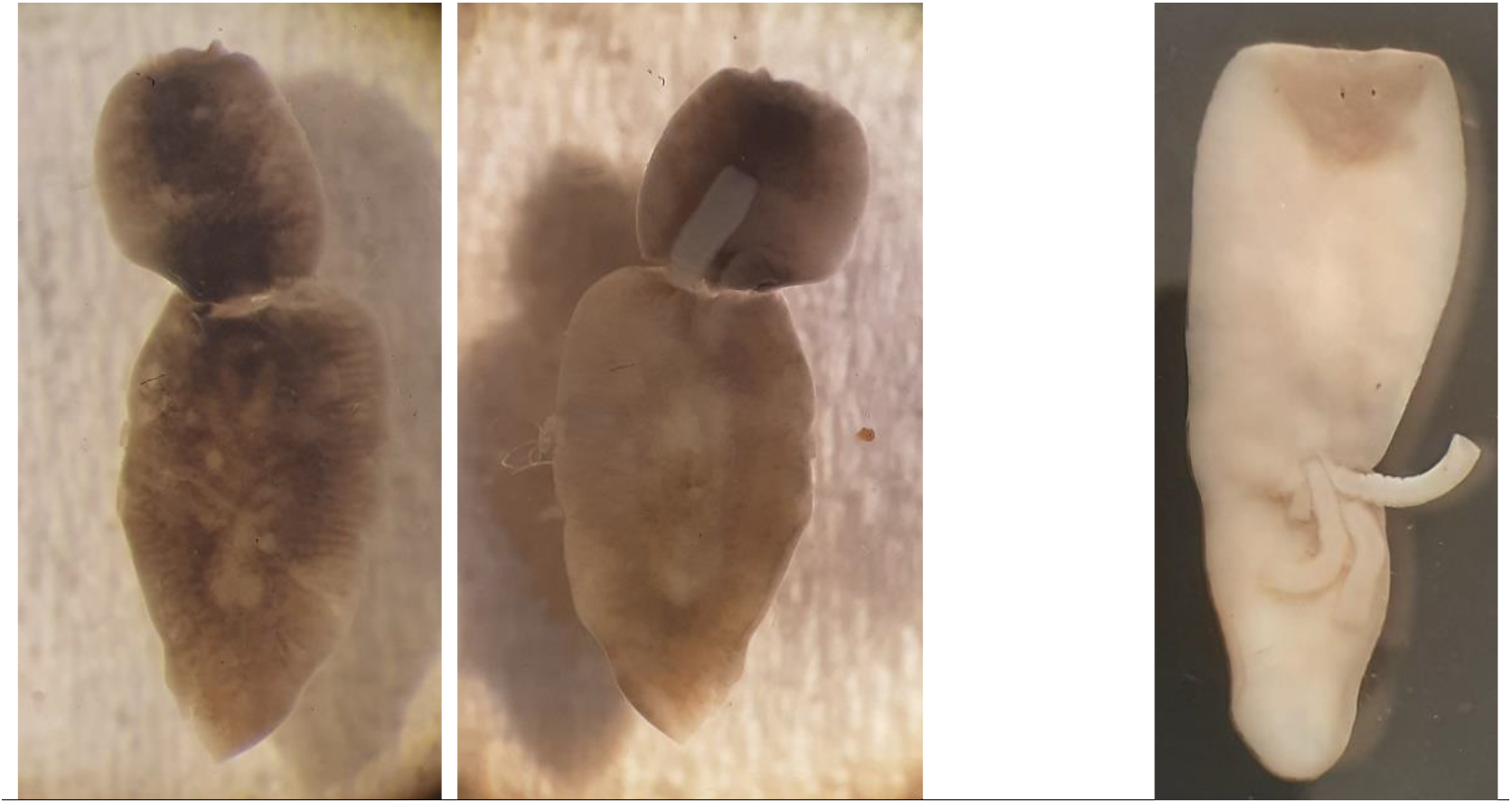
Polypharyngeals undergoing fission – the left group of 2 pictures: in November; right picture: in July

**Fig. 22.**
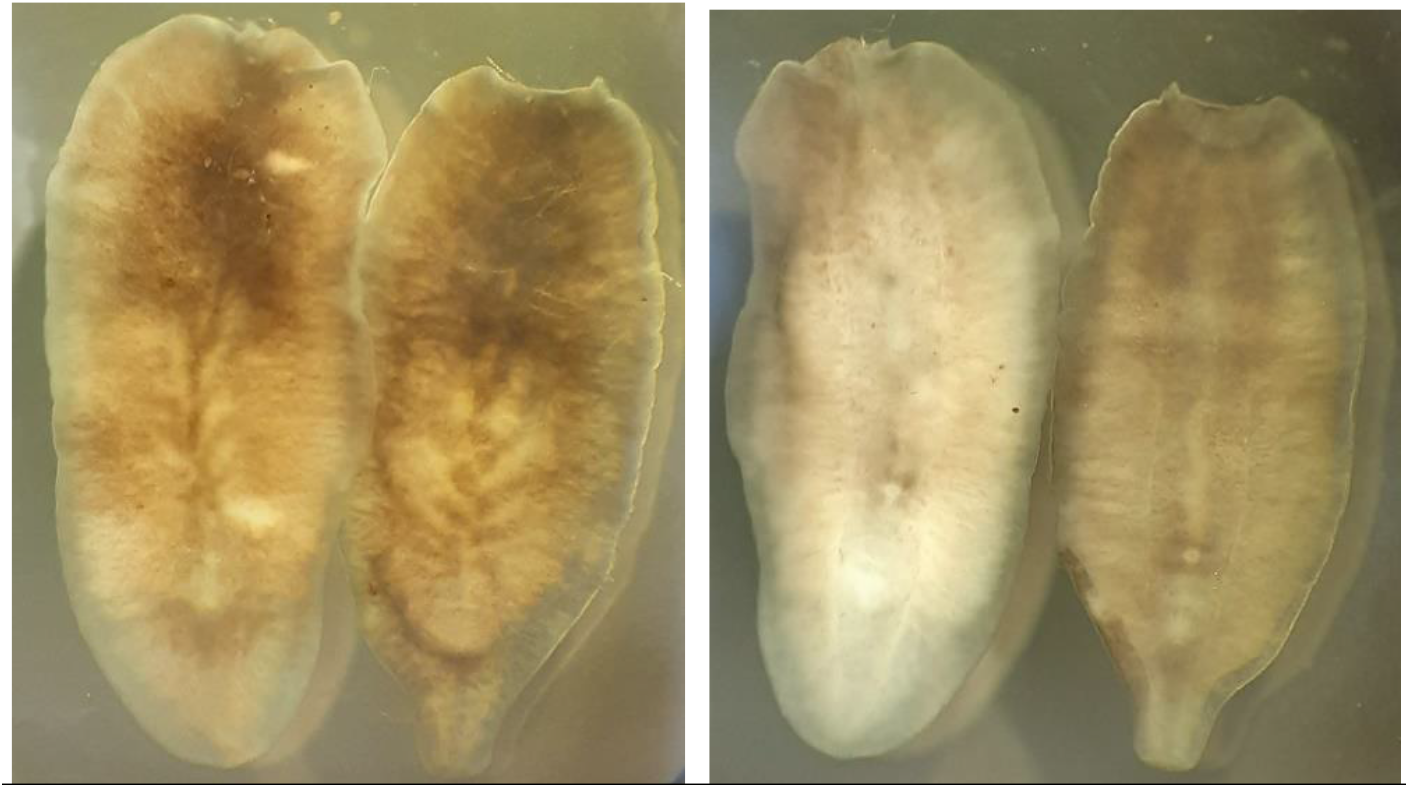
Polypharyngeal specimen (the specimen on the right) probably resulted from fission, 7.11 2022, Rânca, swampy area, spring to the triple fountain, the rear end slightly tapering should be noted

No internal or external cocoons have been observed.

Mating was not observed in either form.

In resume, a putative and incomplete biological cycle can be sketched. Although I did not observed mating, I assume that they mate. Nevertheless, auto-fecundation should not be ecluded.

The **monopharyngeals** mate and produce internal cocoons during the warm season (June, July). The few asexuals or with small copulatory area are most probably resulted (may be resulted) from cocoons hatching. These hatchlings grow and start to develop the copulatory area. They become specimens with small copulatory area, unable for sperm transfer in November. The November asexuals (with no copulatory area) result from late summer hatching or from fission.

The **polypharyngeals** mate during the cold season, they have large copulatory area. They are unable to mate during the warm season, they have small copulatory area or lack it.

This summary strongly suggests the reproductive isolation of the two forms (the monopharyngeal *alpina* and the polypharyngeal *montenigrina*) in mixed populations. The reproductive isolation is a seasonal one, the two forms become capable for mating in different seasons.

## 4. Discussions

### 4.1 Morphological comparison with other populations of *Crenobia alpina* and *Crenobia montenigrina* (in Sluys 2022)

From the analysis of Table 2, I conclude that the monopharyngeal Crenobia subject of this paper belongs to *Crenobia alpina* considered as separate species by Sluys (2022).

**Table 2:**
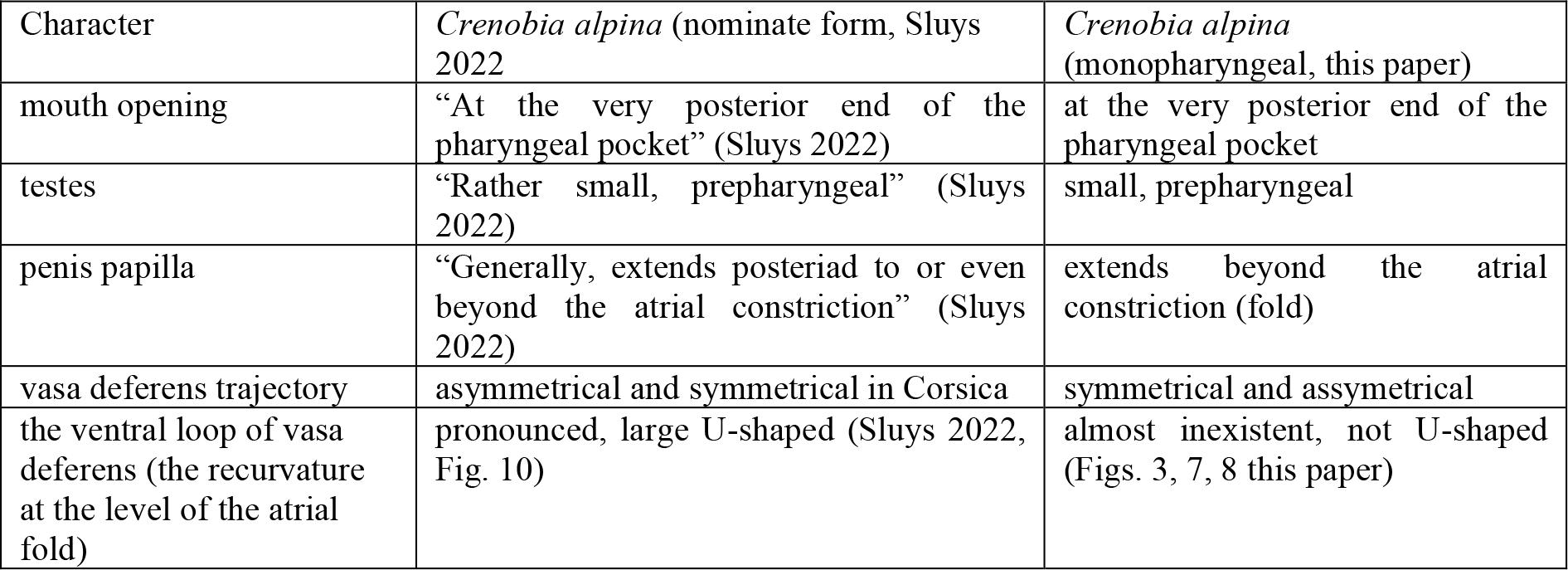
Crenobia alpina.

From the analysis of Table 3, I conclude that the polypharyngeal subject of this paper belong to *Crenobia montenigrina* considered a full species by Sluys (2022).

**Table 3:**
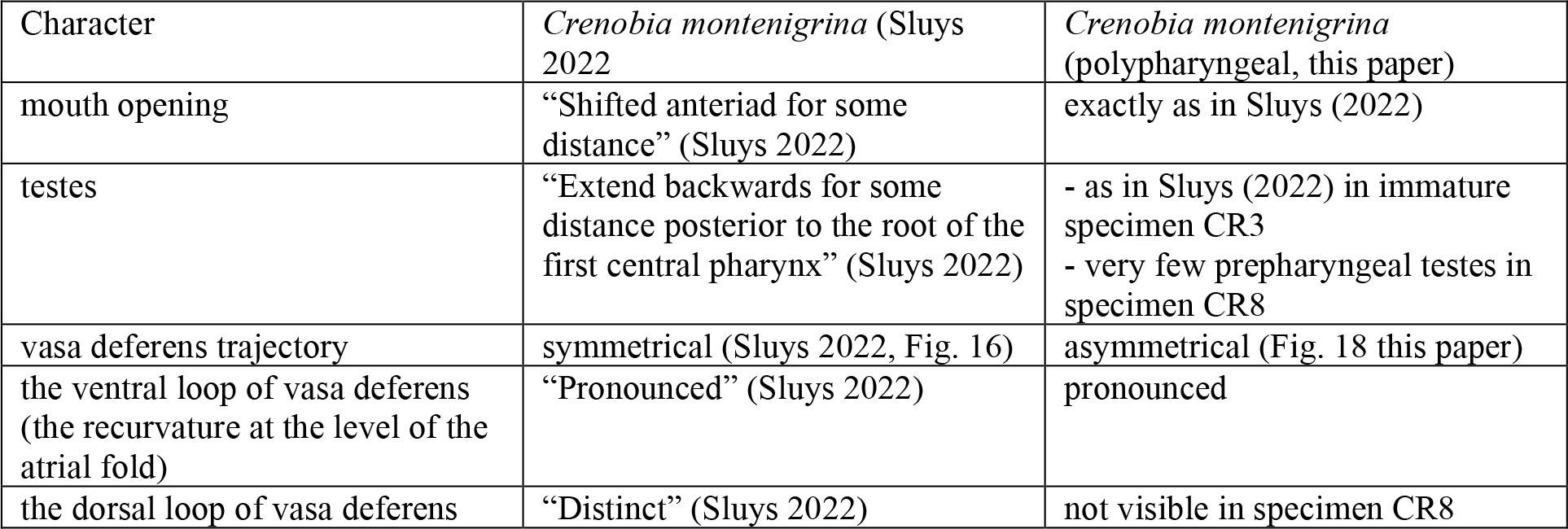
Crenobia montenigrina.

The symmetry of the vasa deferens in specimen CR1 (monopharyngeal) and the asymmetry in specimen CR8 (polypharyngeal) might be an individual character or even a micro-populational level character.

### 4.3 The complex system of fine ducts / sclerotized lines

might represent either artefacts (histological processing defects) or genuine biological structures with unknown functional / physiological meaning. I think for genuine biological structures.

### 4.4 The reproductive characteristics

of *Crenobia alpina* and *Crenobia montenigrina* of this paper differ:

- the polypharyngeals undergo fission during both warm and cold season
- in monopharyngeals, I did not observe the fission

### 4.4 The aspect of the pharynx

in the very small monopharyngeal specimen presented in the right part of Fig. 20 makes me wonder: how does a polypharyngeal hatch – as a monopharyngeal or as a polypharyngeal?

### 4.5 The reproductive isolation

should be verified by the comparative study of spermatozoa in the two forms.

## Appendix

**Fig. 1.**
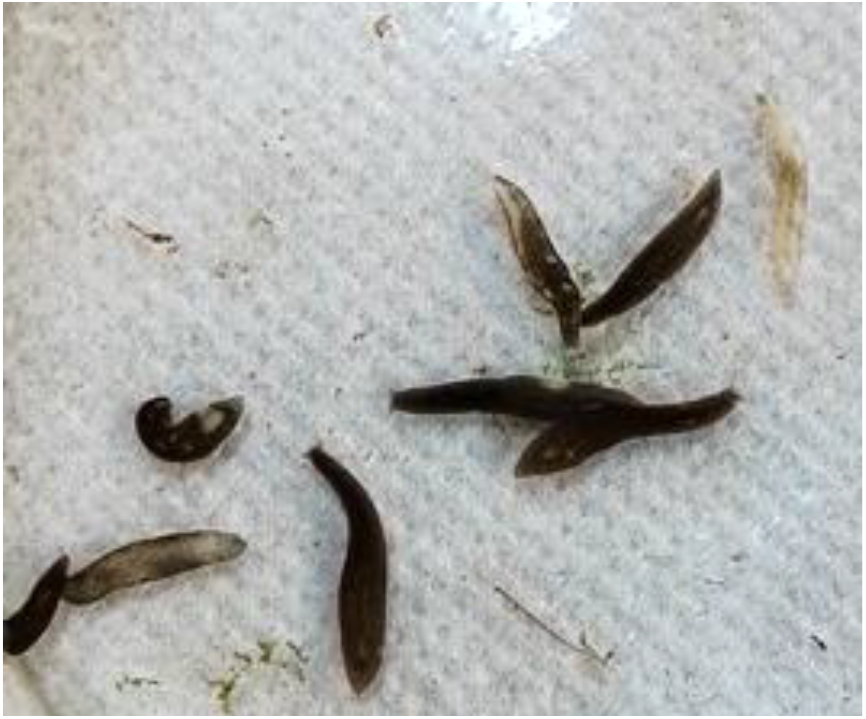
Appendix: unpigmented, black and marbled polypharyngeal specimens

## Notes

### Competing Interest Statement

The authors have declared no competing interest.

